# Spatial transcriptomics reveal pitfalls and opportunities for the detection of rare high-plasticity breast cancer subtypes

**DOI:** 10.1101/2023.04.24.538061

**Authors:** Angèle Coutant, Vincent Cockenpot, Lauriane Muller, Cyril Degletagne, Roxane Pommier, Laurie Tonon, Maude Ardin, Marie-Cécile Michallet, Christophe Caux, Marie Laurent, Anne-Pierre Morel, Pierre Saintigny, Alain Puisieux, Maria Ouzounova, Pierre Martinez

**Affiliations:** Université de Lyon, Université Claude Bernard Lyon 1, INSERM 1052, CNRS 5286 Centre Léon Bérard, Cancer Research Center of Lyon, Lyon, France; Department of Pathology, Centre Léon Bérard, Lyon, France; Centre Léon Bérard, Lyon, France; Plateforme de bioinformatique Gilles Thomas, Synergie Lyon Cancer, Centre Léon Bérard, Lyon, France; Department of Medical Oncology, Centre Léon Bérard, Lyon, France; Institut Curie, PSL Research University, Paris, France; Chemical Biology of Cancer Laboratory, CNRS UMR 3666, INSERM U1143, Paris, France

## Abstract

Breast cancer is one of the most prominent types of cancers, in which therapeutic resistance is still a major clinical hurdle. Specific subtypes like Claudin-low (CL) and metaplastic breast cancers (MpBC) have been associated with high non-genetic plasticity, which can facilitate resistance. The overlaps and differences between these orthogonal subtypes, respectively identified by molecular and histopathological analyses, are however still insufficiently characterised. Adequate methods to identify high-plasticity tumours to better anticipate resistance are furthermore still lacking. Here we analysed 11 triple negative breast tumours, including 3 CL and 4 MpBC samples, *via* high-resolution spatial transcriptomics. We combined pathological annotations and deconvolution approaches to precisely identify tumour spots, on which we performed signature enrichment, differential expression and copy-number analyses. We used the TCGA and CCLE public databases for external validation of expression markers. By levying spatial transcriptomics to focus analyses only to tumour cells in MpBC samples, and therefore bypassing the negative impact of stromal contamination, we could identify specific markers that are not expressed in other subtypes nor stromal cells. Three markers (*BMPER, POPDC3* and *SH3RF3*) could furthermore be validated in external expression databases encompassing bulk tumour material and stroma-free cell lines. We find that existing bulk expression signatures of high-plasticity breast cancers are relevant in mesenchymal transdifferentiated compartments but can be hindered by stromal cell prevalence in tumour samples, negatively impacting their clinical applicability. Spatial transcriptomics analyses can however help identify more specific expression markers, and could thus enhance diagnosis and clinical care of rare high-plasticity breast cancers.

## Introduction

Breast cancer is one of the most prominent types of cancers, with 2.3 million women diagnosed in 2020 (Globocan, 2020 IARC). Patient stratification relies on the presence of specific targetable alterations in the oestrogen receptor (ER), progesterone receptor (PR), and HER2 genes. These genetic properties are generally well recapitulated by the broad PAM50 transcriptomic signatures: luminal A, luminal B, HER2-enriched, basal-like and normal-like^1^. Clinical approaches typically rely on the immunohistochemical expression of those three markers, done routinely during the pathological examination of breast cancer specimens. Triple negative breast cancers (TNBC) patients however lack any of the targetable ER, PR and HER2 alterations, leading to scarce therapeutic options and low survival rates despite promising results with antibody-drug conjugates^2^. Rarer subtypes such as claudin-low (CL) tumours or metaplastic breast carcinoma (MpBC) have furthermore been associated with high plasticity, a cellular property facilitating dynamic phenotypic changes and the subsequent emergence of non-genetic therapeutic resistance^3^. Anticipating breast cancer cells’ ability to adapt *via* plasticity is thus of paramount importance for effective therapeutic targeting. However, the driving mechanisms of tumour plasticity remain poorly understood, and no standard method exists to accurately detect nor quantify it for patient stratification.

High-plasticity breast cancer subtype identification typically relies on either molecular or histopathological analyses. Claudin-low tumours were originally defined by transcriptomic analyses, with a phenotype similar to basal cells lacking expression of claudins 3, 4, and 7, and other cell-cell adhesion markers^4^. They represent 3-5% of all breast cancers^5,6^, and are generally associated with strong stemness features. The evolutionary trajectories explaining their malignant progression are however still debated. CLs typically display high expression of epithelial-mesenchymal transition (EMT) factors ^4^, known to foster phenotypic plasticity and stemness^7,8^, but also enhanced tolerance to oncogenic stress, thereby mitigating genomic instability^9^. Recent work further suggests the existence of different claudin-low classes: CL1, CL2 and CL3^5^. CL1 tumours are believed to arise directly from malignantly transformed mammary stem cells (MaSC). These tumours were also the most similar to previous observations of EMT-driven plasticity and genomic stability, displaying the highest intrinsic expression of EMT markers and the lowest fraction of genome altered (FGA).

Metaplastic breast carcinomas (MpBC) are a heterogeneous group histopathologically-defined by the presence of a non-epithelial tumour component, believed to occur through transdifferentiation^10^. MpBCs account for 0.2-2% of all invasive breast carcinomas are usually triple negative and often associated with poorer survival rates^11^. Different sub-types exist according to the transdifferentiated component^11^, including but not restricted to spindle, chondroid and osseous cells^11,12^. MpBC can present one or more metaplastic component, which are often admixed with a component of invasive breast carcinoma of no special type (IBC-NST). MpBC diagnosis remains challenging and adequate markers are still lacking to correctly classify specific subtypes within this highly heterogeneous disease^13^. Similarly to CLs, many of these sub-types display high EMT marker expression and resemble mammary tumour–initiating stem cell–like cells, based on transcriptomic data^14^. Although CL tumours frequently involve metaplastic differentiation^4^, the full extent of the overlaps and divergences between these subtypes defined by different approaches is still unclear.

Here, we aimed to further characterise these plasticity-associated breast cancer subtypes, defined either molecularly or histopathologically, *via* spatial transcriptomics (SpaT). Based on previously described^5,6^ genomic instability and CL-associated bulk gene signatures, we identified 3 putative CL tumours (CL-like) and 4 non-CL, genomically unstable TNBC samples as controls. We also selected 4 MpBCs through histopathology (2 spindle cell, 1 chondroid, 1 IBC-NST compartment from a mixed spindle cell tumour), and performed SpaT analyses on all 11 samples. We report that, unlike in unstable TNBC samples, the tumour compartment of CL-like tumours did not recapitulate expression patterns expected from bulk analyses. We find that existing CL expression signatures are significantly upregulated in MpBCs with mesenchymal trandifferentiation, but that the prevalence of stromal cells can hinder clinical applicability and lead to false positive diagnoses. This pitfall highlights the need to integrate histopathological approaches into transcriptomics analyses to define more robust signatures that are specific to high-plasticity tumour cells only.

By levying SpaT to focus expression analyses only to tumour cells in MpBC samples, we could identify specific expression markers that are not expressed in other subtypes nor in stromal cells. SpaT-derived markers could thus enhance the diagnosis and clinical care of rare high-plasticity breast cancers in the future.

## Material and Methods

### Low and high genomic instability sample identification

From a larger cohort of 379 samples with RNA-seq data assembled as part of the MyPROBE project (17-RHUS-0008), we analysed 87 fresh frozen TNBC samples with paired whole-exome sequencing (WES) data collected at Centre Léon Bérard. Only samples with more than 30% tumour purity (estimated percentage of tumour cells), as estimated by FACETS^15^, had been included in this cohort. We used the already processed RNA-seq and WES data available for all samples. Using the pre-calculated fractions of genome altered (FGA), we identified 3 samples with low genomic instability (FGA <10%) and 6 samples with high instability (FGA>75%). Samples CLB-17, CLB-52 and CLB-74 were considered CL-like, samples CLB-14, CLB-23, CLB-37, and CLB-51 were considered unstable TNBC (Supplementary Table 1). An additional CL-like sample (CLB-11, CL-like 4) was discarded after in-depth investigation identified an erroneously low FGA (actual FGA > 10%). A tumour-free slide from this discarded tumour was however included in the controls.

### Spatial transcriptomics analyses

10x Genomics Visium (Spatial 3’ v1 for fresh frozen samples) slides were processed according to the manufacturer’s guidelines and sequenced in two batches on a NovaSeq 6000 Illumina sequencer, targeting 50,000 reads/spot: A first batch included samples M1 to M11 (Cl-like, unstable TNBC, normal controls), a second batch included samples M13 to M16 (metaplastic TNBCs). The spaceranger software was used to process the raw data. Stereoscope^16^ was used for deconvolution analyses using only the TNBC data from a single-cell breast cancer atlas^17^, and excluding plasmablasts that were initially found to be over-represented in deconvolution results (included cell types: cancer epithelial, normal epithelial, endothelial, CAF, T-cell, B-cell, myeloid, PVL).

Signature enrichment analyses were performed using the AUCell tool, designed for UMI-based single-cell data that present similar biases to those of our SpaT data. To maximise the capture of lowly-expressed genes in all signatures, we used the maximum threshold advised by AUCell developpers (0.2). This provided higher minimal scores compared to lower thresholds (0.5, 0.1, 0.15), without altering the overall observations (Supplementary Fig. 1).

### Combining histopathological annotations and spatial transcriptomics on fresh frozen tissue

We analysed a total of 15 Visium capture areas, including 3 controls extracted from tumour samples with epithelial compartments but no tumour identifiable on the slide. All slides were annotated by a breast pathologist, to separate tumour/non-tumour and epithelial/non-epithelial compartments, as well as eliminate spots corresponding to folded tissue and artefacts. The SpaT data and paired annotations have been publicly deposited as a series on the Gene Expression Omnibus (GSE213688).

Tumour spots were first annotated by the pathologist. To later investigate expression markers that would not be biased by the prevalence of mesenchymal stromal cells, we restricted the stromal spots definition to the following pathologist annotations, exclusively: Fibrosis, Fibrosis (peritumoral), Fibrous stroma, Tumour Stroma, Tumour stroma fibrous, the last two separated on visual appreciation of the cellular density and collagen abundance in the stroma. Tumour and stromal spots were further refined using scores obtained by deconvolution, to keep only the spots most enriched in the population of interest. Tumour spots with a cancer epithelial score < 0.1 were discarded. In addition, tumour spots in the MpBC_chondroid sample with unusual B-cell scores > 0.1 were also discarded. For stromal spots, those with either normal epithelial or cancer epithelial scores > 0.1 were discarded.

### CNA profiles

Three slides that were devoid of tumour cells upon histopathological examination, were included as normal references: M1 and M9 (CL-like3) and M7 (CL-like4). The expression profiles of epithelial spots present on these slides provided a more relevant reference than those of other cells, in which cell-type-specific expression patterns local to genome segments could hamper CNA identification. Individual spot copy number (CN) profiles were produced with the infercnvPlus tool (https://github.com/CharleneZ95/infercnvPlus, based on inferCNV of the Trinity CTAT Project: https://github.com/broadinstitute/inferCNV). All spots annotated as either tumoral or epithelial from every slide were pooled together, using the epithelial ones as references.

This produced a relative CN measure per gene per spot, which we summarised into a major cytoband per spot matrix, by computing the average CN per cytoband. Sample-specific profiles were then obtained by averaging the relative CN measures per cytoband across all spots of each given sample. Comparable profiles were obtained for bulk samples, by averaging per cytoband the segmented logR values reported by FACETS^15^ analysis, from the whole exome sequencing data of CL-like and unstable samples.

Gain/loss profiles were calculated for each bulk sample by attributing values of 1 (gain), 0 (normal) or -1 (loss) to each cytoband, according to whether their mean logR exceeded the median logR of the sample ± 0.6, as implemented in ^18^. A threshold was used to calculate similar gains/loss profiles in Visium samples, according to whether the cytoband CNA scores exceeded the threshold. Its value was optimised to minimise the pairwise distance across all cytobands between the bulk-derived and SpaT-derived gain/loss profiles for the 4 unstable tumours, which provided the most reliable and informative data.

### Differential expression analyses on SpaT data

All tumour samples were first individually normalised using the SCTransform function from the Seurat R package^19^. All 11 samples were then merged together and re-normalised using the SCTransform function, resulting in a matrix of 22,058 genes by 14,905 spots. The MAST R package^20^ was used to performed differential expression analyses on SpaT data. Fold changes (FC) were defined as the mean expression in the population of interest divided by the mean expression in the control population.

The CL, unstable control and normal samples (M1 to M11) were distributed over 3 Visium slides and were sequenced together in a first batch. All 4 MpBC samples (M13 to M16) were on the same Visium slide and were sequenced in a second separate batch. We thus adapted our design to remove potential biases stemming from batch effects and different sequencing depths. The UMAP projection of all MpBC tumour spots was obtained without spots from other samples, to prevent batch effects, then normalised using the harmony software^21^ to account for technical effects between capture areas. For the same reasons, we initially defined differentially expressed (DE) genes only among the metastatic samples. We used the tumour spots from the MpBC_NST sample as a non-metastatic reference, and the spots annotated as fibrosis and tumour stroma from the MpBC_spindle2 sample as a stromal reference. To account for the imbalance in number of tumour spots between the two spindle MpBC samples, we first performed DE analysis between them, and retained all genes whose expression was not significantly different (p > 0.05 and absolute log2 FC < 2). We then performed DE analyses between the union of tumour spots from both spindle samples against the tumour spots from the NST sample and the stromal references, separately. Genes upregulated in the spindle tumour spots were defined as those having a log2 FC > 1.5 and a corrected p-value < 0.001 (Benjamini-Hochberg). Genes overexpressed in MpBC tumour spots in both analyses (vs non-MpBC tumour and stroma) were considered as candidate MpBC markers. We further validated their relevance in the tumour and stromal spots from the CL-like and unstable samples, using the same method but with a higher log2 FC threshold (>2) in this bigger dataset. Only genes significantly overexpressed in MpBC samples in both analyses (non-MpBC tumour and stroma) were considered as internally validated markers. For the external validation, we used the clinical information, including sample metaplastic status, and mRNA expression (z-normalised RNA-seq log RSEM) data from the TCGA breast cancer Firehose Legacy cohort, retrieved through cBioPortal^22,23^. CCLE data, z-normalised relative to diploid samples, was similarly obtained from cBioPortal. CL classification of the CCLE breast cancer cell lines was retrieved from ^5,24^. Differential expression was assessed using Wilcoxon rank-sum tests for each gene of interest.

## Results

### Identification of CL-like, unstable and metaplastic breast tumours

We decided to focus on identifying CL tumours displaying high intrinsic plasticity and low genomic instability, as described by the CL1 sub-classification. We analysed 379 TNBC samples from Centre Léon Bérard for which RNA-seq data was available, 87 of which also had paired whole-exome data. To first identify potential CL1 tumours based on genomic stability, we selected 3 tumours with FGA < 10%. For control TNBC samples, we decided to focus on those least resembling CL1 tumours and identified 6 genomically unstable tumours with FGA > 75% (Supplementary Table 1). The low-FGA tumours displayed RNA expression patterns highly concordant with CL, and particularly CL1, phenotypes: high *ZEB1, ZEB2*, and *MSRB3*; low *POLQ*^9,24^ (Fig. 1A). We thus considered these 3 samples as highly relevant CL-like candidates. Out of the 6 high-FGA samples, to be used as controls, we selected the 4 high-FGA samples displaying the most contrasting patterns regarding these genes (Supplementary Fig. 2). We performed PAM50 centroid classification on the 379 samples from our cohort merged with the TCGA BRCA dataset. This revealed that the 3 CL-like samples were considered as Luminal A, and the 4 unstable control samples were considered as Basal. These findings were furthermore consistent with gene-set enrichment analyses of established CL and Basal-like gene signatures from ^4^, and the CL1 and MaSC signatures from ^5^ (Figure 1B-C, Supplementary Fig. 3-5). These tumours were histologically reviewed to confirm their histotype, all were classified as triple negative breast carcinoma of no special type (NST) according to the WHO Classification of Breast Tumours, 5^th^ ed. criteria, and no metaplastic component was identified in these samples.

**Figure 1:**
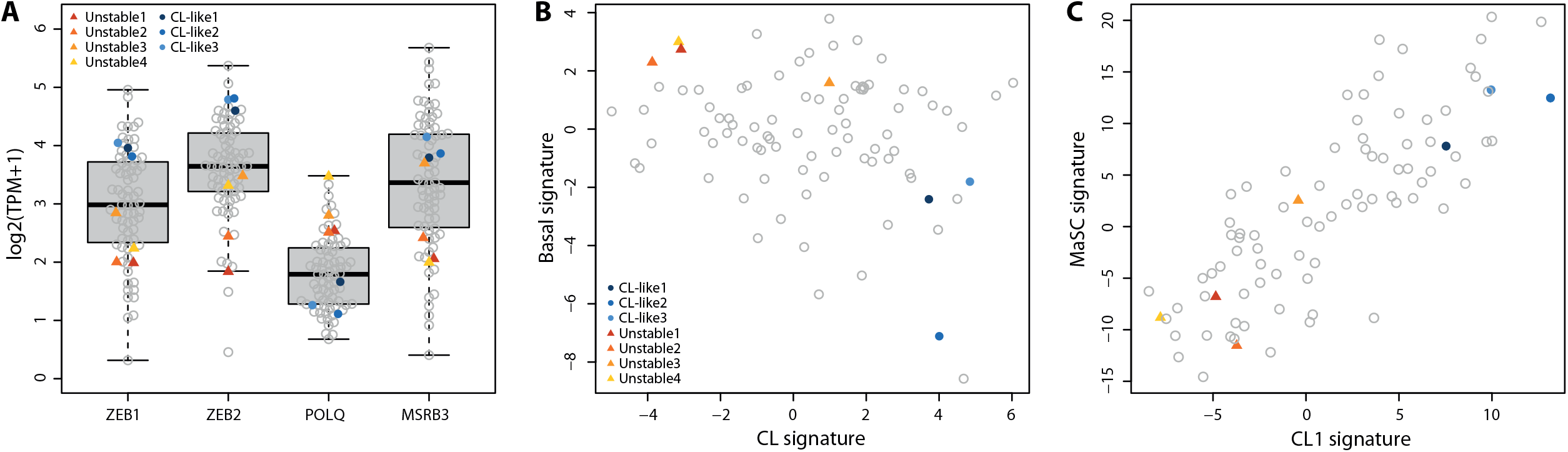
Identification of CL-like and unstable samples. Expression of CL1 markers *ZEB1, ZEB2, POLQ* and *MSRB3* in the 87 MyPROBE TNBC samples with paired WES and RNA-seq data for (A) 3 low-FGA samples considered as CL-like and (B) 4 high-FGA samples considered as unstable. Scores for gene-set enrichment analyses across the 87 samples for (C) the Claudin-low (CL) and Basal expression signatures and D) the Claudin-low type 1 (CL1) and Mammary Stem Cell (MaSC) expression signatures.

To broaden the analysis focusing on high-plasticity breast tumour types, we further selected 4 MpBC, of different subtypes: 2 spindle cell carcinomas (MpBC_spindle), 1 carcinoma with pure chondroid differentiation (MpBC_chondroid) and 1 IBC-NST compartment from a tumour diagnosed as a mixed spindle cell and IBC-NST (MpBC_NST). Subtypes were initially determined during routine histopathological evaluation for clinical diagnosis on FFPE samples, and were reviewed afterward by a breast pathologist. Only tumours for which fresh frozen material was available were selected, in order to use the same spatial transcriptomics technology as for CL and control TNBCs. Frozen specimens were also reviewed on hematoxylin and eosin stained (H&E) slides, to assess the sampled components and select optimal samples to be used for spatial transcriptomic techniques.

### Spatial transcriptomics data deconvolution reflects histopathological annotations

We performed spatial transcriptomics (SpaT) analyses on a total of 14 slides comprising: 3 CL-like tumours; 4 unstable TNBCs; 4 MpBCs; and 3 adjacent normal tissue with no detectable tumour cells. We report medians of 85% reads under tissue, 2,398 genes per spot and 5,083 UMIs per spot (Supplementary Table 1). Estimation of spot cellular composition using the stereoscope deconvolution software confirmed that spots histopathologically identified as tumour displayed more tumour-associated expression patterns (p < 0.001, t-test, Figure 2, Supplementary Fig. 6-10), and that adjacent normal slides were tumour-free (Supplementary Fig. 9). This confirms that the RNA profiles from tumour-labelled spots reflected pathological annotations.

**Figure 2:**
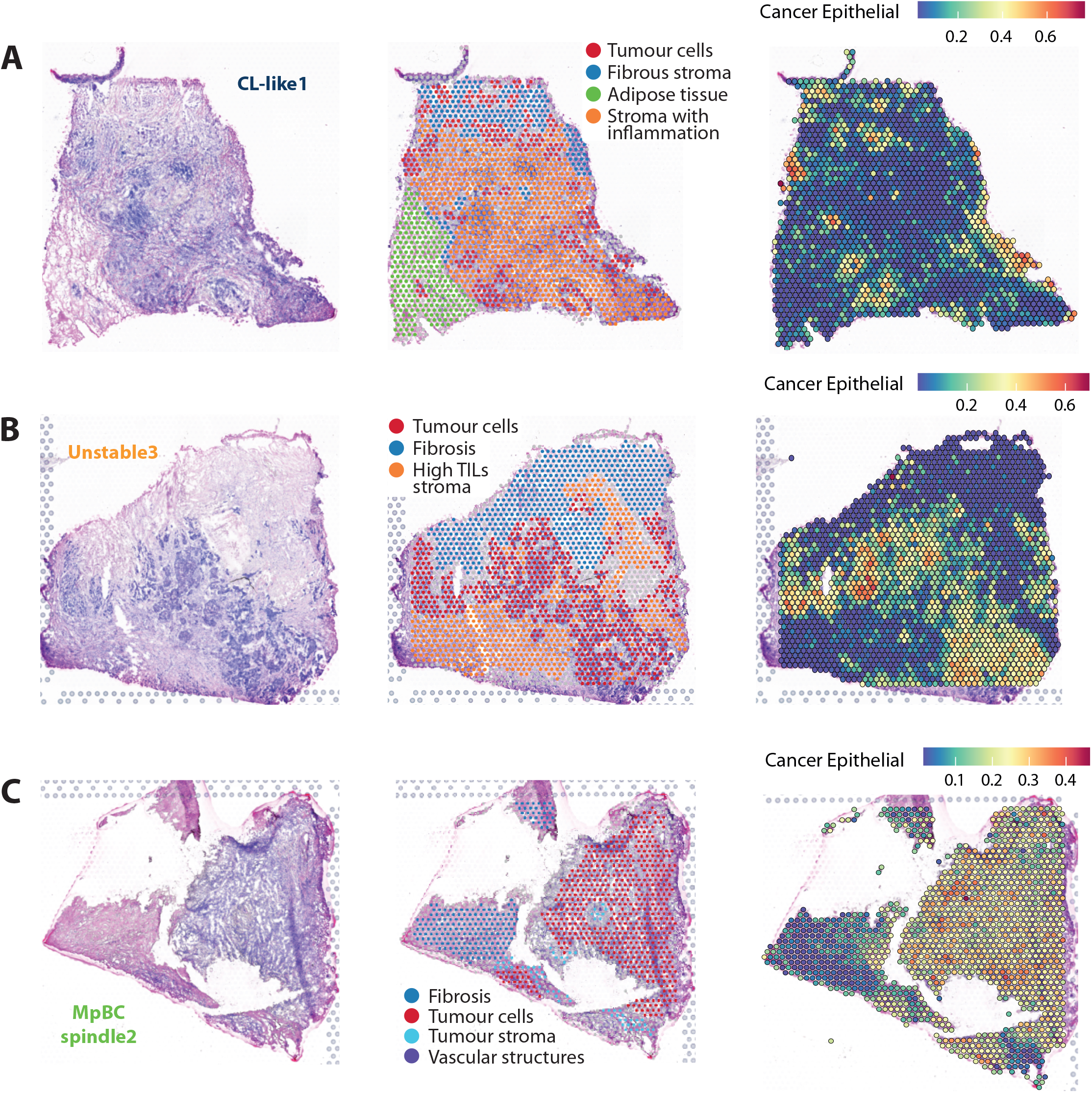
Overlaying pathology annotations and in-silico deconvolution in SpaT data. Hematoxylin & eosin staining (left), pathologist annotations (centre) and per-spot deconvolution-based cancer epithelial score (right) for samples (A) CL-like1 (B) Unstable3 and (C) MpBC_spindle1. Unlabelled grey spots were either considered artefacts or could not be annotated with confidence by the pathologist.

Automated deconvolution was based on individual RNA profiles of cells extracted from tumours of epithelial origin^17^, but could identify cancer cells in all but one of the 4 MpBC slides: the MpBC_spindle1 sample is the only exception (Supplementary Fig. 8), in which the strong mesenchymal nature of metaplastic tumour cells led to spots being predicted to mostly contain cancer-associated fibroblasts. In later analyses, we furthermore used both pathological annotations and thresholds on deconvolution scores to more rigorously identify genuine tumour and non-tumour spots (see Methods). Of note, all SpaT analyses were performed on tumour sections that were distinct from the ones initially used for bulk analyses, which can lead to sampling bias. Importantly, we report that the tumour slides from the CL-like2 and CL-like3 analysed by SpaT were annotated as non-invasive in situ carcinoma, while both patients were diagnosed with invasive TNBC. This may imply phenotypic differences compared to the sections included in bulk analyses initially performed on these samples.

### Spatialisation sheds light on the micro-environmental impact on molecular plasticity signatures

To assess how SpaT recapitulate the bulk signal used to classify the CL-like and unstable control samples, we generated pseudo-bulk data by individually pooling all spots together for each slide. In addition, histopathological annotations of tumour spots allowed us to investigate the prominence of CL, Basal, CL1 and MaSC signatures on the entire surface of slides, but also specifically in tumour spots. When including all spots per slide, correlations between the bulk and pseudo-bulk (all spots) data were statistically significant for the Basal and CL1 signatures (rho=0.93, p=0.007; rho=0.82, p=0.034, respectively), but not for the CL and MaSC signatures (Fig. 3A). However, no correlations were found to be significant when focusing solely on tumour spots (Figure 3B). This suggests that focusing on tumour spots only provides a worse recollection of the signal initially used to determine CL status in bulk analyses.

**Figure 3:**
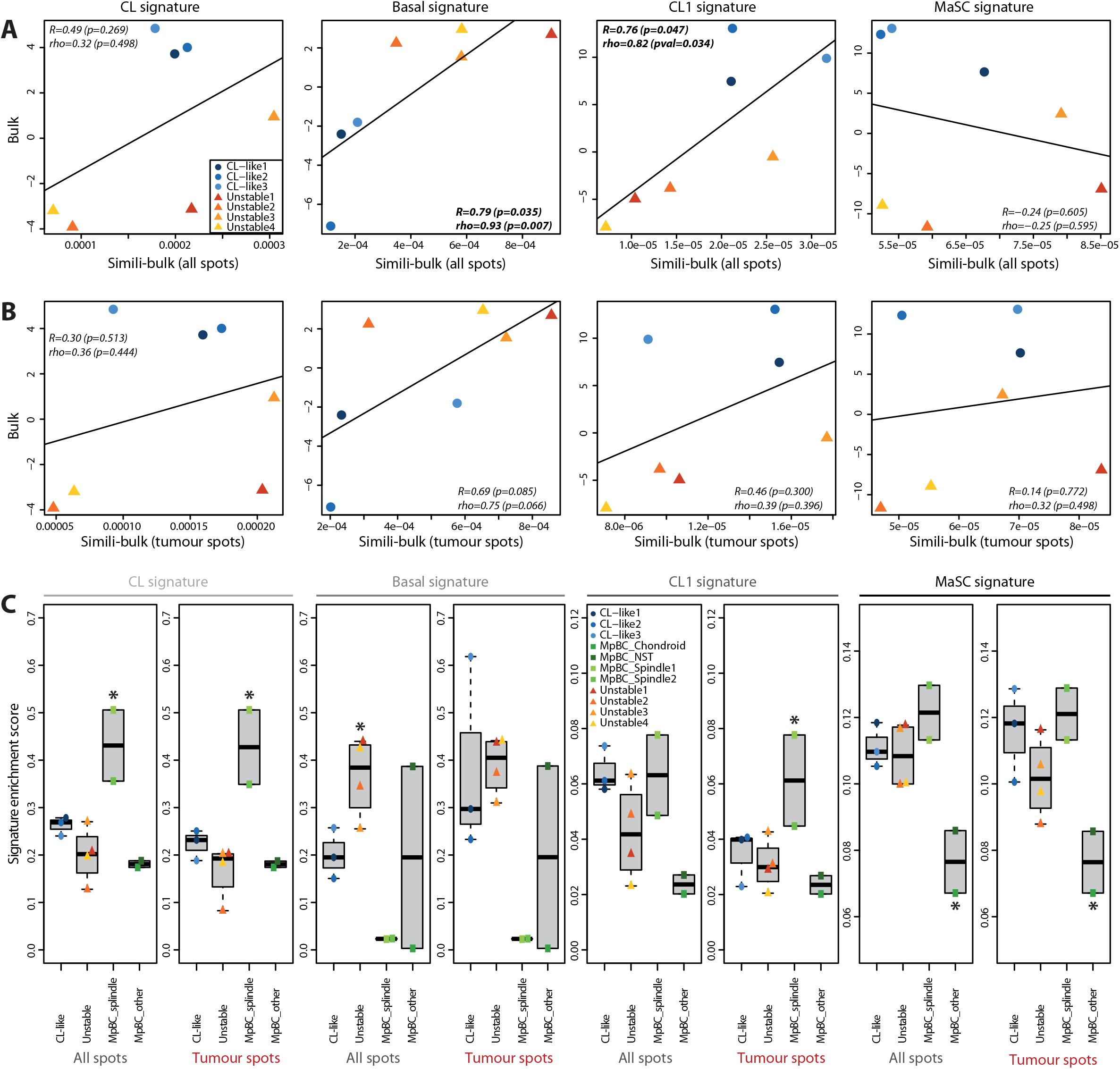
Spatialised enrichment analyses of existing plasticity-associated expression signatures. (A) Correlation for gene-set enrichment scores between bulk and SpaT-derived pseudo-bulk data for the CL, Basal, CL1 and MaSC expression signatures, using all annotated spots. (B) Correlation for gene-set enrichment scores between bulk and SpaT-derived pseudo-bulk data for the CL, Basal, CL1 and MaSC expression signatures, focusing on tumour spots only. (C) Average per-spot gene-set enrichment scores for the CL, Basal, CL1 and MaSC expression signatures, in each individual sample of the CL-like, unstable and MpBC sample types. For clarity, the MpBC samples were dichotomised according to the presence or not of a spindle-cell transdifferentiated compartment on the captured area (“spindle” or “other”, respectively). * highlight groups of samples whose signature enrichment scores were significantly different from all other samples pooled together (p<0.05, Wilcoxon rank-sum tests).

We further analysed each signature specifically, using either all spots or only tumour ones in all samples (Figure 3C). The original CL signature was significantly higher in spindle cell MpBC samples, particularly in tumour spots only (Figure 3C, p=0.036, Wilcoxon rank-sum tests). The CL1 signature was also enriched in spindle cell MpBC tumour spots (p=0.036). The 3 spindle cell and chondroid MpBC samples also displayed significantly lower enrichment for the Basal signature, when grouped together (Figure 3C, p=0.012, both cases). This suggests that these existing signatures are suited to the identification of breast cancer transdifferentiation, particularly to a mesenchymal phenotype.

The restriction of signature enrichment analyses to tumour spots had little impact on unstable control samples. In CL-like samples however, CL and CL1 signatures decreased in this case, while the Basal one increased, albeit only with borderline significance due to the few data points (p=0.169, p=0.020 and p=0.254, respectively, t-tests). This further indicates that the bulk-based signal used to identify CL-like samples was not reflected by the specific transcriptomic profiles of tumour cells in these samples.

### CL-like samples poorly recapitulate bulk copy number alterations

We determined relative copy number (CN) profiles per major cytoband for each SpaT tumour spot, using the InferCnvPlus software (Fig. 4A). Spot-specific profiles were then averaged to obtain per-sample CN profiles (Supplementary Fig. 11A). These profiles derived from measured RNA quantities are only relevant as relative values within each sample and thus cannot be used to determine absolute copy numbers. We however defined a threshold to identify regions of chromosomal loss and gain based on the unstable control samples, which offered more reliable expectations in terms of number CNA per sample and signal quality (Supplementary Fig. 11B-C, Supplementary Fig. 12-14, see Methods). This allowed us to calculate relative fractions of genome altered (rFGA), and to investigate distances and correlations between bulk and SpaT CN profiles (Fig. 4B-C). Contrarily to expectations from bulk data, CL-like samples did not appear genomically stable and their SpaT-derived rFGAs was not significantly different from those of unstable control samples (p=0.88, t-test). We however found that CL-like CN profiles appeared less correlated with their bulk counterparts than unstable profiles, both on the entire genome (Supplementary Fig. 15) and exclusively on regions of chromosomal gain or loss (p=0.064 and p=0.074, respectively, t-test). Analyses of differences in rFGA and average pairwise distances between bulk and SpaT profiles (see Methods) furthermore confirmed that the bulk CNA information was much better recapitulated by SpaT data in unstable control samples than in CL-like samples (Fig. 4C-D, both p<0.001, t-test).

**Figure 4:**
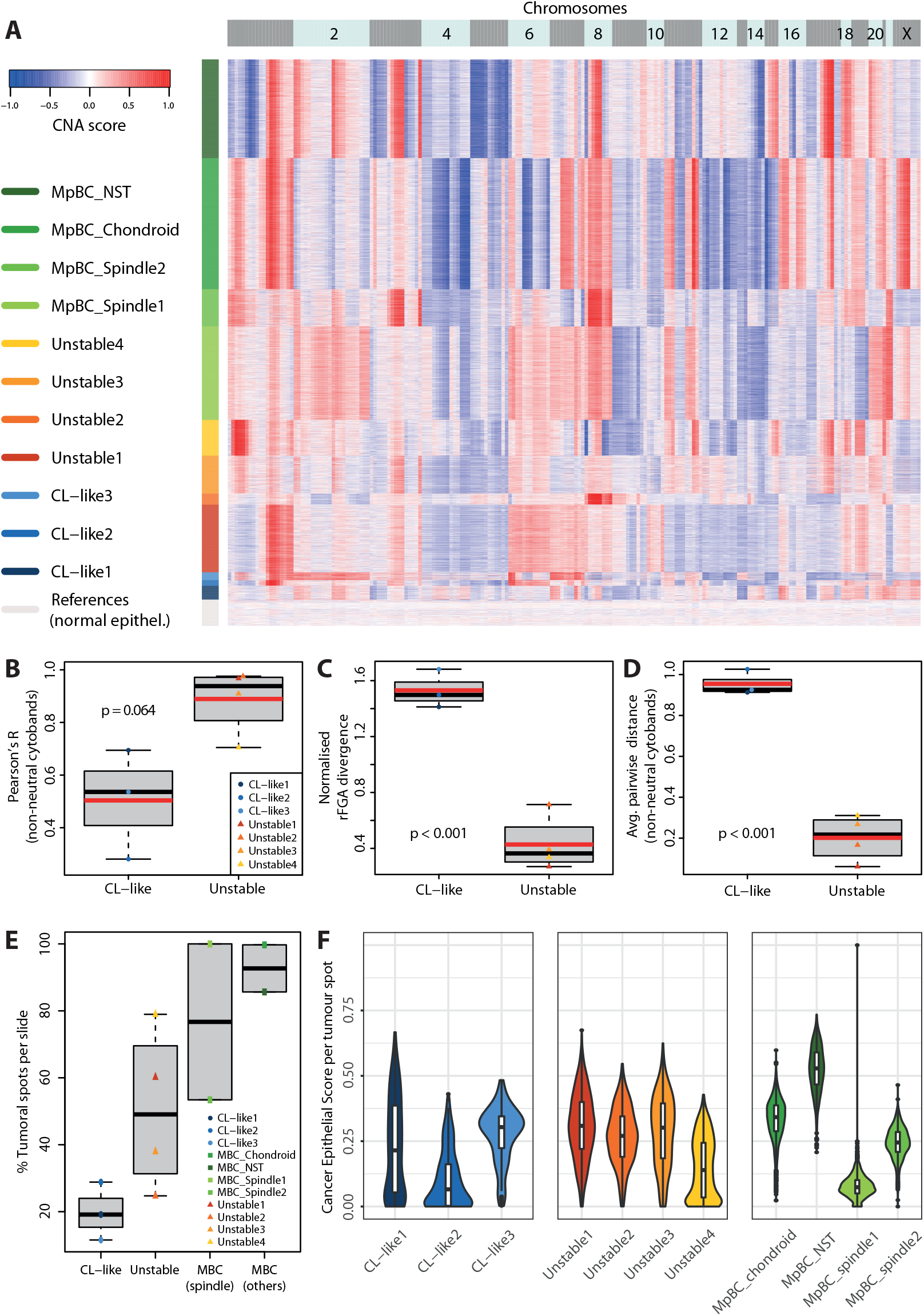
Unstable SpaT-derived copy-number profiles are less divergent from their bulk counterpart. (A) Copy-number alterations (CNA) profile calculated for each tumour spot of all cancer 11 samples. This score represents a per-spot relative measure, with values close to -1 (blue) corresponding to the highest loss of genomic material, and values close to 1 to the highest gain. For each spot, values were averaged per major cytoband, *i*.*e*. the average of the scores obtained for each individual gene in the cytoband. (B) Pearson correlation between bulk logR and SpaT-derived average CNA scores, averaged per major cytoband, in CL-like and unstable samples. Only cytobands harbouring non-neutral CNAs in the SpaT data were included in the correlation calculation. (C) Normalised divergence between bulk- and SpaT-derived relative Fractions of Genome Altered, in CL-like and unstable samples. (D) Average pairwise distances between bulk- and SpaT-derived copy number profiles in CL-like and unstable samples. CN profiles were based on a -1 (loss), 0 (neutral) or 1 (gain) attributed to each cytoband. Only the cytobands reported as non-neutral in the SpaT profiles were included in the distance calculation. (E) Percentage of tumour spots in each captured area, per sample type. (F) Cancer epithelial score, as reported by stereoscope-based deconvolution, for all tumour spots in each sample.

### Higher stromal content in CL-like samples

We report that the SpaT slides from CL-like tumours contained a significantly smaller fraction of tumour spots (Fig. 5E, p=0.02, Wilcoxon rank-sum test), and a lower fraction of cancer epithelial cells per spot than the other tumours, as estimated by stereoscope, even including transdifferentiated MpBC samples (p < 0.001, Wilcoxon rank-sum test). Given that tumour spots in CL-like samples did not display strong signal for the highly mesenchymal CL expression signatures, this strongly suggests that an insufficient percentage of tumour cells in CL-like bulk samples could have both artificially increased enrichment scores for claudin low expression signatures, and hampered the detection of genuine CNAs.

**Figure 5:**
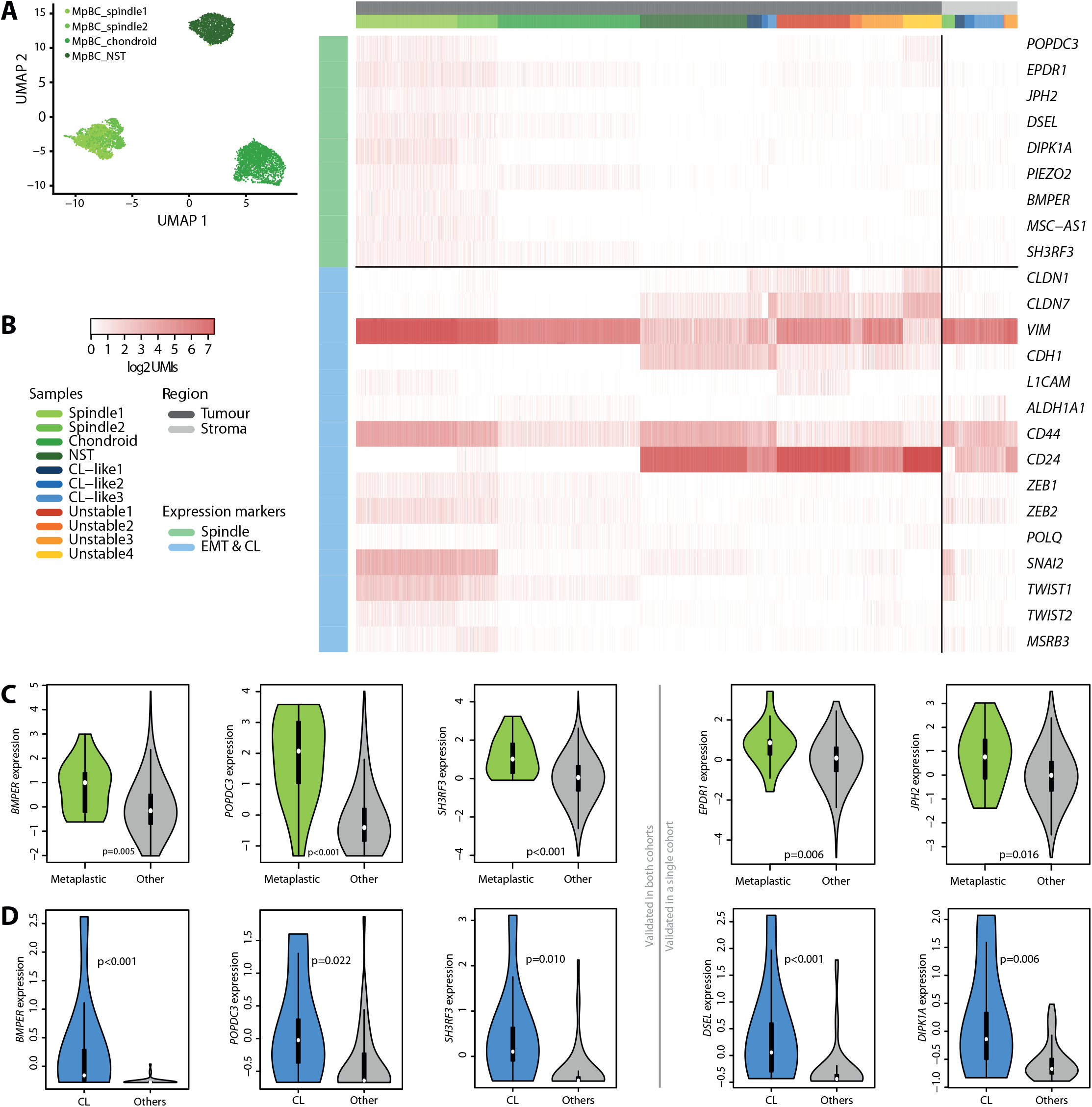
SpaT-derived expression markers. (A) UMAP projection of all tumour spots from the 4 MpBC samples analysed by SpaT. (B) Expression heatmap for the newly identified spindle cell MpBC markers (top, light green) and known EMT and CL markers (bottom, light blue). Expression is reported in log2 number of UMIs per spot, ranging from white (no expression) to red (high expression), and was analysed in both tumour spots (left, black) and stromal spots (right, grey). (C) Expression of validated spindle cell MpBC genes *BMPER, EPDR1, JPH2, POPDC3* and *SH3RF3*, respectively, in an external cohort of 1108 samples analysed per bulk RNA-seq, stratified by tumour metaplastic status (14 metaplastic tumours in total). (D) Expression of validated spindle cell MpBC genes *BMPER, POPDC3, SH3RF3, DSEL* and *DIPK1A*, respectively, in an external cohort of 51 cancer cell lines analysed per RNA-seq, stratified by CL status (9 CL samples in total).

### SpaT can identify tumour-specific markers that are robust to stromal content

Clustering analyses on the tumour spots from MpBC samples revealed that both spindle cell samples clustered together, highlighting that they shared common, spindle-cell specific expression patterns (Fig. 5A). We thus harnessed SpaT data to identify genes overexpressed in MpBC tumour cells compared to both tumour cells of no special type and healthy stromal cells.

With 1,983 spots per slide on average (range: 1,055 - 3,037), our SpaT data can provide a very powerful basis for differential expression analyses, even with few samples. We however aimed to mitigate the impact of the low number of samples and maximise reproducibility (see Methods). We first took advantage of the high number of spots available, and split our cohort into internal discovery (MpBC samples) and validation (CL-like and unstable samples) sets to prevent overfitting (Table 1). We identified genes significantly overexpressed in spindle cell or chondroid spots compared to both NST and stroma spots from MpBC samples, combining two separate differential expression analyses. We then selected only those also overexpressed compared to both tumour spots and stromal spots from CL-like and unstable control samples.

**Table 1:**
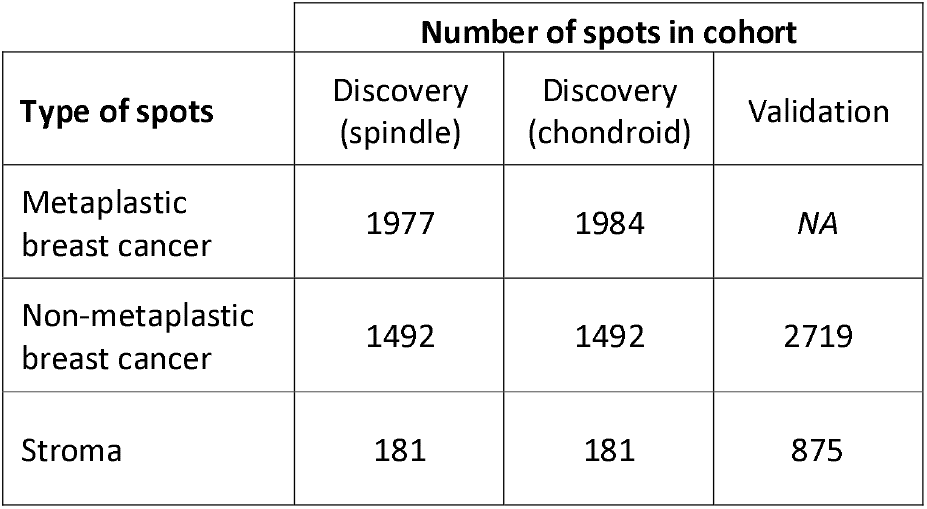
Number of spots used in each design for differential expression analyses. Metaplastic breast cancer spots were compared to both non-metaplastic breast cancer spots and stroma spots separately, first in the discovery cohort, then in the validation cohort.

Using this design, we could identify subtype-specific genes with low-expression in non-metaplastic tumour cells and stromal cells. 9 genes were overexpressed in the two spindle cell MpBC samples: *BMPER, DIPK1A, DSEL, EPDR1, JPH2, PIEZO2, POPDC3, MSC-AS1* and *SH3RF3* (Fig. 5B). We also investigated known EMT markers^4–6^, none of which came up as significant in our analyses, and whose expression patterns were less specific to spindle cell MpBC tumour cells. The same analysis conducted on the MpBC_chondroid samples identified 8 genes whose overexpression was specific to chondroid MpBC cells: *HSPB6, VGLL3, PTX3, GFRA1, MT1G, MT1E, CHI3L2* and *SAA1* (Supplementary Fig. 16).

These genes are however identified from a small number of samples (n=2 for spindle cell, n=1 for chondroid). To assess their relevance in larger cohorts, we investigated their expression in two complementary external datasets: the TCGA breast cancer expression dataset comprising 1108 samples^25^, including 14 metaplastic ones; and the 51 stroma-free breast cancer cell lines from the CCLE^26^, including 9 samples classified as Claudin-low. Long non-coding RNA *MSC-AS1* was the only gene for which expression data was unavailable, in both sets. In the TCGA dataset, samples were classified as non-CL (n=940), CL1 (n=69), CL2 (n=42) or CL3 (n=57) according to previous work based on expression signatures^24^. *BMPER, EPDR1, JPH2, POPDC3* and *SH3RF3* were significantly overexpressed in metaplastic samples (all p<0.05, Wilcoxon rank sum tests; Fig. 5C, Supplementary Fig. 17). In the CCLE dataset, *BMPER, DIPK1A, DSEL, POPDC3* and *SH3RF3* were significantly overexpressed in CL cell lines (Fig. 5D, Supplementary Fig. 18). All significant p-values held after Benjamini-Hochberg correction in each dataset.

These results validate that 3 out of the 8 analysable stroma-independent spindle cell MpBC markers (*BMPER, POPDC3* and *SH3RF3*) are highly relevant in metaplastic tumours and CL cell lines from two independent external cohorts. Four more genes could furthermore be validated in one external cohort (*EPDR1* and *JPH2* in the TCGA data; *DIPK1A* and *DSEL* in the CCLE data), but not the other. These results suggest that leveraging SpaT data to focus solely on tumour cells can yield powerful expression markers that are less likely to be biased by stromal cell prevalence. This can help identify high-plasticity tumours more reliably, and improve molecular classification.

In addition, all genes except *PIEZO2* furthermore displayed significant overexpression in CL subtypes (Supplementary Fig. 19) in the TCGA data, confirming the similarities between CL and MpBC tumours. The overlap between the two subtypes showed significant enrichment, as 57% of TCGA (8 / 14) metaplastic breast cancers were classified as CL (4 CL1, 1 CL2 and 3 CL3), which represent only 15% of tumours (p<0.001, Fisher’s exact test). As for chondroid markers, genes *VGLL3* and *PTX3* could both be validated in metaplastic breast cancer samples in the TCGA data, due to their significant overexpression after multiple testing (Supplementary Fig. 20).

## Discussion

Rare breast cancer subtypes associated with cellular plasticity features are still difficult to both diagnose and treat. In particular, there appears to be an overlap between molecularly-defined claudin-low (CL) tumours and metaplastic breast cancers (MpBC) typically defined by histopathology. Here we harnessed the novel possibilities offered by recent advances in spatial transcriptomics (SpaT) to shed light on the commonalities and discrepancies between these plasticity-associated breast cancer subtypes, and assess the opportunities and pitfalls their routine classification presents. We identified 3 putative CL samples and 4 unstable TNBCs using gene expression and copy number data, as well as 4 MpBCs reviewed by a breast pathologist. We analysed a single slide of each of the 11 samples by spatial transcriptomics, along with 3 slides from adjacent normal breast tissue with no detectable tumour material as controls.

We investigated 4 previously reported signatures (Claudin-low *vs* Basal from ^4^, Claudin-low type 1 and Mammary stem cells from ^5^) and found they were highly relevant in MpBC, particularly for the spindle cell sub-classification. We however found that the CL-like samples, defined by molecular analyses, were heavily impacted by stromal cell prevalence. This can hamper the detection of genuine CNAs and bias RNA signal towards mesenchymal gene overexpression, which is a feature of CL tumours. Restricting our SpaT analyses to the spots harbouring tumour cells revealed that these cells did not recapitulate the low-FGA, high CL expression features that prompted our initial assessment of these samples as putative CL, based on bulk analyses.

Overall, these findings illustrate the difficulties hindering molecular-based identification of high-plasticity subtypes, as over-representation of stromal cells in a sample can lead to false positives. This is likely the case for the 3 tumours we identified as putative CL tumours. It is clear that at least a subset of tumours molecularly classified as CL would also be histologically defined as spindle-cell MpBCs. The nature of CL tumours that are not spindle-cell MpBCs remains however insufficiently characterised. It will be important to determine whether these samples merely display strong stromal prevalence, or whether they reflect an additional specific tumoral phenotype. If indeed such cells displaying CL transcriptomics features but without being histologically identified as metaplasia exist, our results here highlight that SpaT analyses could both validate their existence and identify more specific markers. This would help refine the classification of rare breast cancers, and more clearly define non-overlapping *bona fide* subtypes with specific clinical outcomes.

Here we could furthermore harness the spatial information to analyse differential expression between the different compartments of tumour samples at near single-cell resolution. Using internal and external validation procedures, we could identify *BMPER, POPDC3* and *SH3RF3* as robust spindle-cell MpBC markers, whose detection is unlikely to be affected by the stromal content of samples. Of note, *BMPER* has been reported to promote invasive phenotypes and angiogenesis in cancer^27^, and *SH3RF3* has been reported to promote stem-like properties in breast cancer^28^. We furthermore identified *MSC-AS1*, a long non-coding RNA that has been linked to both osteogenic differentiation^29^ and oncogenesis^30^, as an additional potential spindle-cell MpBC markers. We could however not fully validate its relevance due to its absence in the external dataset.

Our SpaT analyses revealed that although breast tumour cells displaying CL-like mesenchymal properties could be detected in spindle cell MpBC samples, their classification using existing molecular signatures in bulk samples remains error-prone. However, even given our limited sample size, SpaT proved extremely powerful for the identification of genes that are highly specific to transdifferentiated tumour cells. This is of particular interest for translational research on rare subtypes, as large cohorts are difficult to obtain. Larger SpaT studies than ours could furthermore provide yet more specific expression signatures for MpBC and CL tumour cells. This can prove useful to help diagnose such complex cases, for which integrative histological and molecular approaches are required to overcome their respective limitations.

## Declarations

### Ethics approval and consent to participate

For the BRC of Centre Léon Bérard (n°BB-0033-00050) biological material collection and retention activity is declared to the Ministry of Research (DC-2008-99 and AC-2019-3426). Samples were used in the context of patient diagnosis. This study was approved by the ethical review board of Centre Léon Bérard. The BRC of Centre Léon Bérard is quality certified according to AFNOR NFS96900 (N° 2009/35884.2) and ISO 9001 (Certification N° 2013/56348.2) for clinical trials, ensuring scientific rigor for sample conservation, traceability and quality, as well as ethical rules observance and defined rules for transferring samples for research purposes (ministry of health for activities authorization n° AC-2019-3426 and DC-2008-99). The use of samples are reviewed by a multidisciplinary committee before any transfer. The samples are properly codified, so that in no case the recipients are able to identify the donor’s identity, or any clinical information that may be used for the donor’s identification. The material used in the study has been collected in agreement with all applicable laws, rules, and requests of French and European government authorities, including the patient’s informed consents.

### Consent for publication

Not applicable

### Availability of data and materials

Spatial transcriptomics data have been deposited as a series on the Gene Expression Ominbus database: GSE213688. The RNA and whole-exome sequencing data from the MyPROBE cohort, analysed to identify the CL-like and unstable samples in this study, have not been published nor released yet. The identifiers will however be consistent with those used here, and data will be accessible upon request to the authors until publication.

### Authors’ contributions

PS, APM, AP, MO and PM designed the study; ML, MCM and CC collected the samples; VC, LM, CD and MO processed the samples; VC annotated the slides; AC, RP, LT, MA and PM performed computational analyses; AC, VC, MO and PM wrote the manuscript.

## Acknowledgements

The biological samples used in this study have been prepared by BB-0033-00050, CRB Centre Léon Bérard, Lyon France.

