## Supplementary Figures and Tables for "Spatial transcriptomics reveal pitfalls and opportunities for the detection of rare high-plasticity breast cancer subtypes"

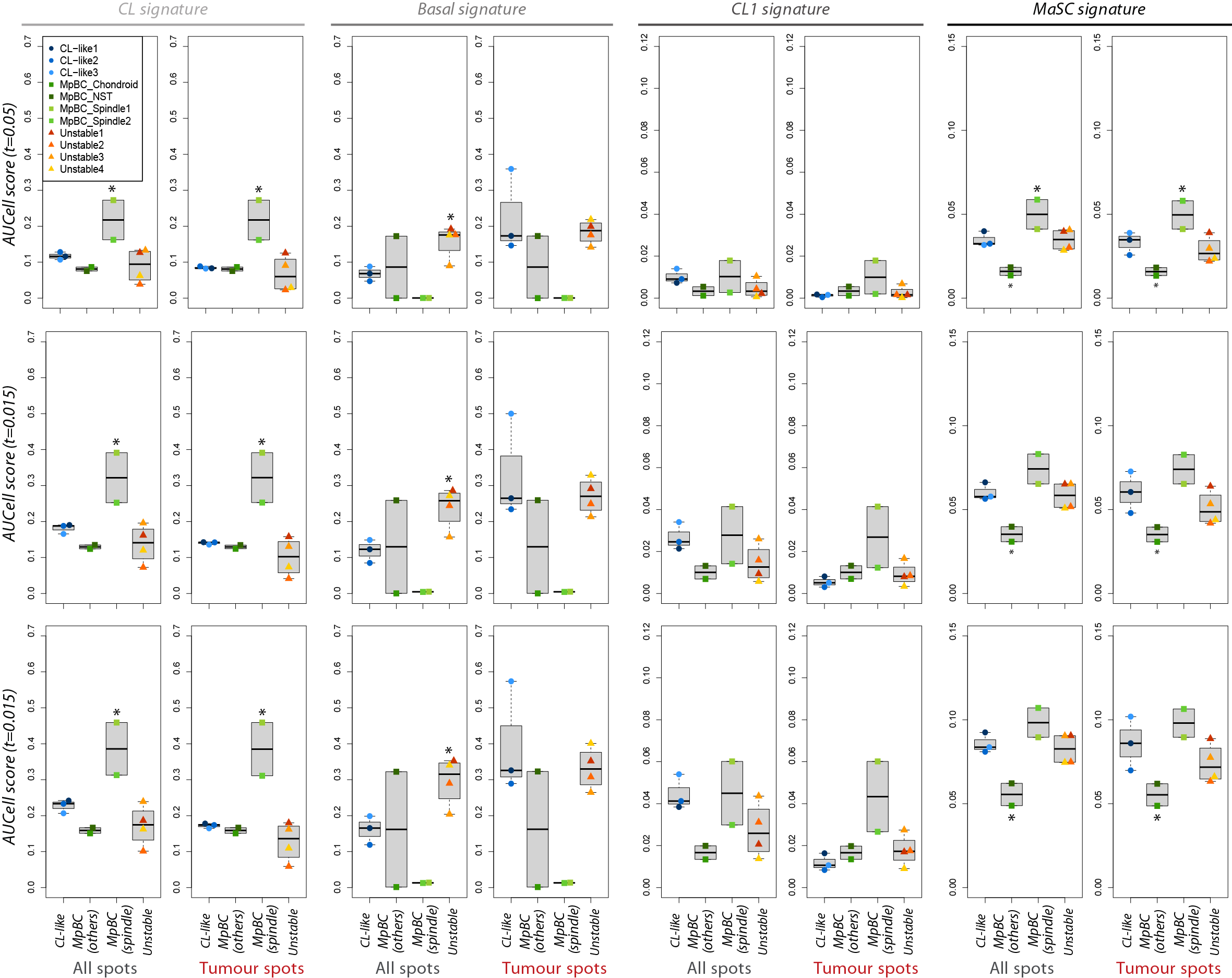

***Supplementary Fig. 1:*** *Impact of AUC threshold on signature enrichment scores.* *Average per-spot gene-set enrichment scores for the CL, Basal, CL1 and MaSC expression signatures, in each individual sample of the CL-like, unstable and MpBC sample types. Different thresholds were used to determine if individual gene are overexpressed in each spot/sample (0.05, 0.10 and 0.15, top to bottom). For clarity, the MpBC samples were dichotomised according to the presence or not of a spindle-cell transdifferentiated compartment on the captured area (“spindle” or “other”, respectively). * indicate groups of samples whose expression if significantly different from all others in each category (p<0.05, Wilcoxon rank-sum test).*

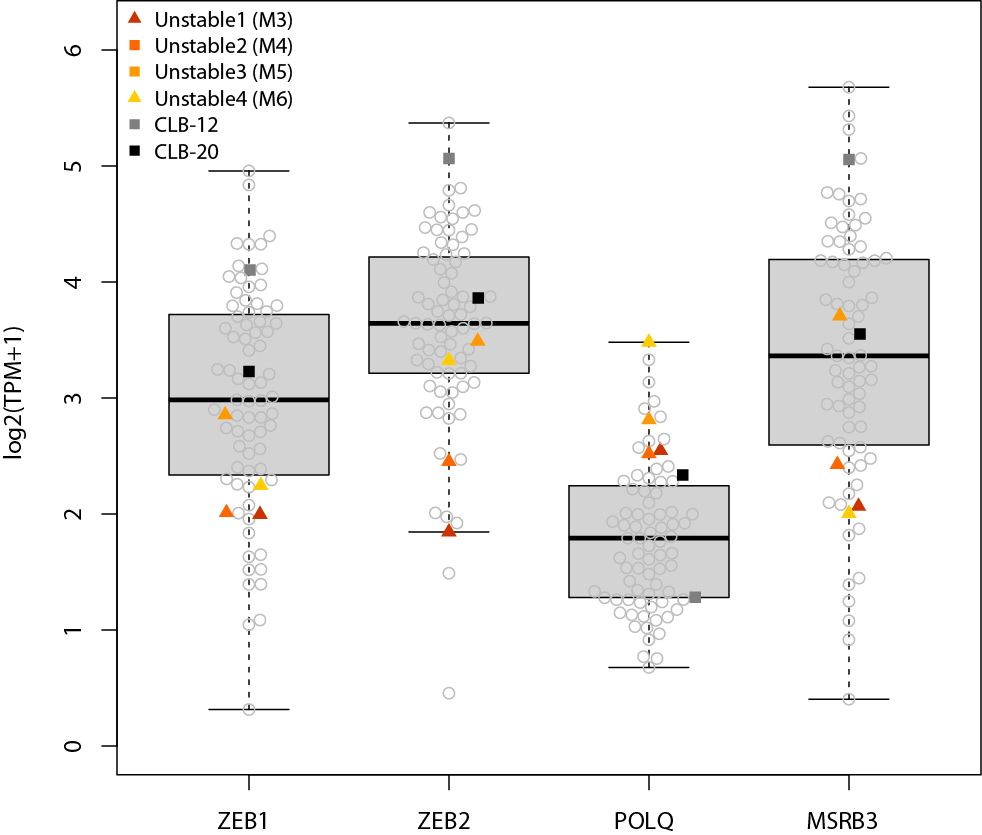

***Supplementary Fig. 2****: Expression of CL1 markers ZEB1, ZEB2, POLQ and MSRB3 in the 87 MyPROBE TNBC samples with WES and RNA-seq data in 6 high-FGA samples. The samples identified by grey and black squares were not considered valid Unstable samples.*

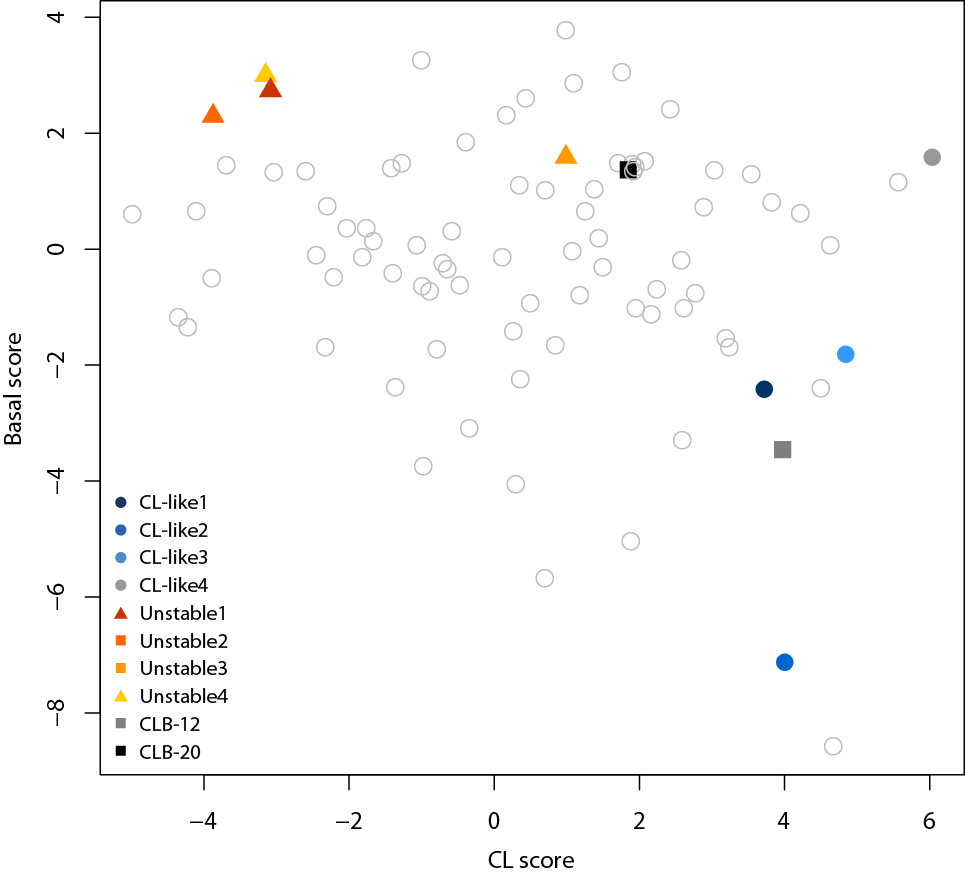

***Supplementary Fig. 3****: Scores for gene-set enrichment analyses across the 87 samples with paired RNA-seq and WES data for the Claudin-low (CL) and Basal expression signatures expression signatures. The high-FGA samples identified by grey and black squares were not considered valid Unstable samples. The CL-like4 sample was later discarded from analyses due to higher FGA than first reported.*

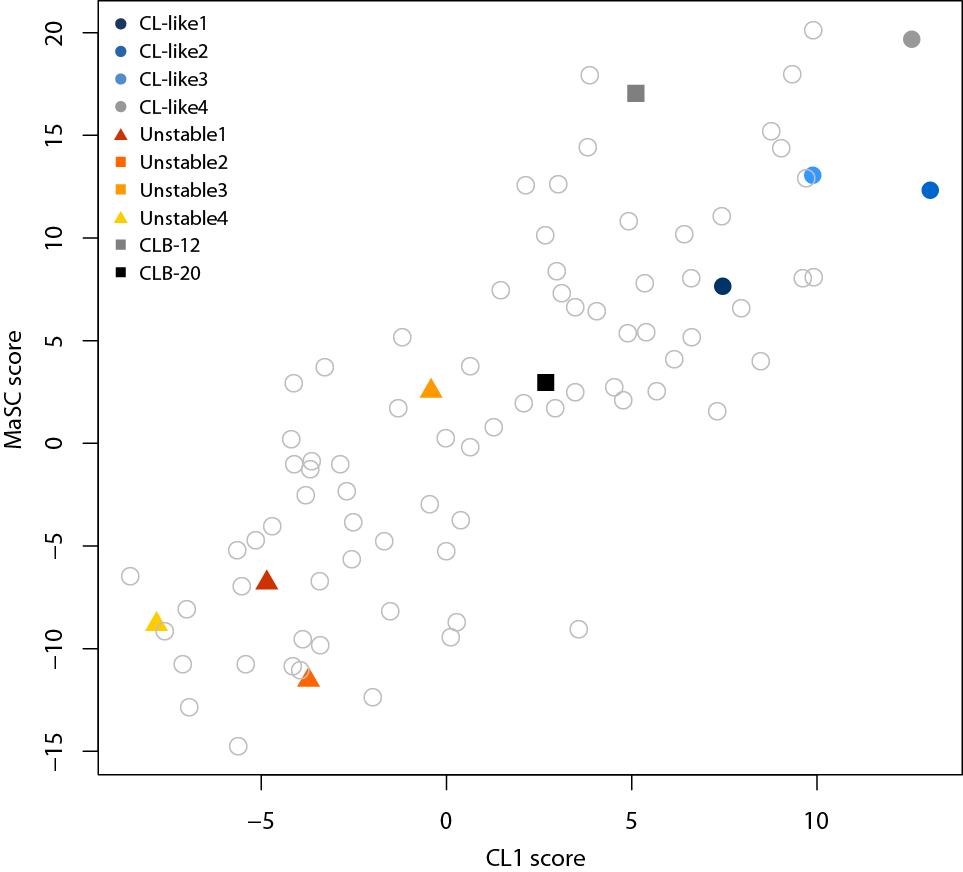

***Supplementary Fig. 4****: Scores for gene-set enrichment analyses across the 87 samples with paired RNA-seq and WES data for the CL1 and MaSC expression signatures. The high-FGA samples identified by grey and black squares were not considered valid Unstable samples. The CL-like4 sample was later discarded from analyses due to higher FGA than first reported.*

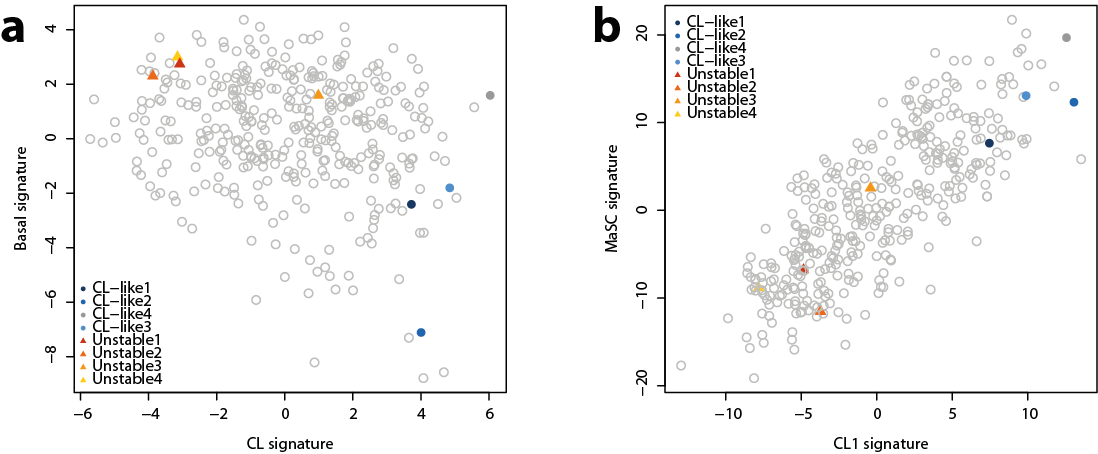
***Supplementary Fig. 5****: Scores for gene-set enrichment analyses across all 379 samples with RNA-seq data for a) the CL1 and MaSC and expression signatures b) MaSC and CL1 expression signatures. The 87 samples with paired WES data are included. The CL-like4 sample was later discarded from analyses due to higher FGA than first reported.*

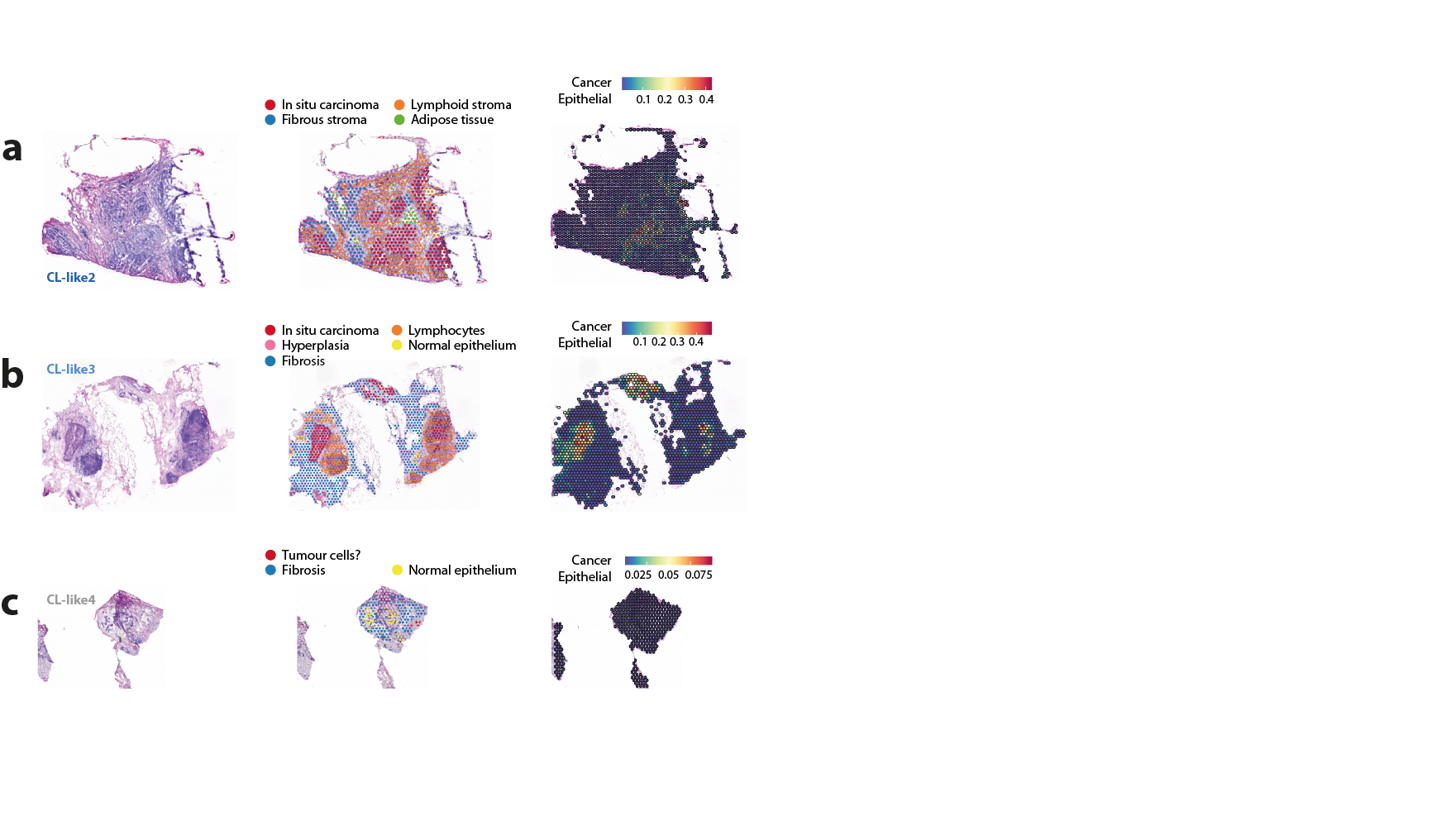

***Supplementary Fig. 6:*** *Hematoxylin & eosin staining (left), pathologist annotations (centre) and per-spot deconvolution-based cancer epithelial score (right) for CL-like samples CL-like2, CL-like3, and CL-like4. The CL-like4 sample was later discarded from analyses due to higher FGA than first reported.*

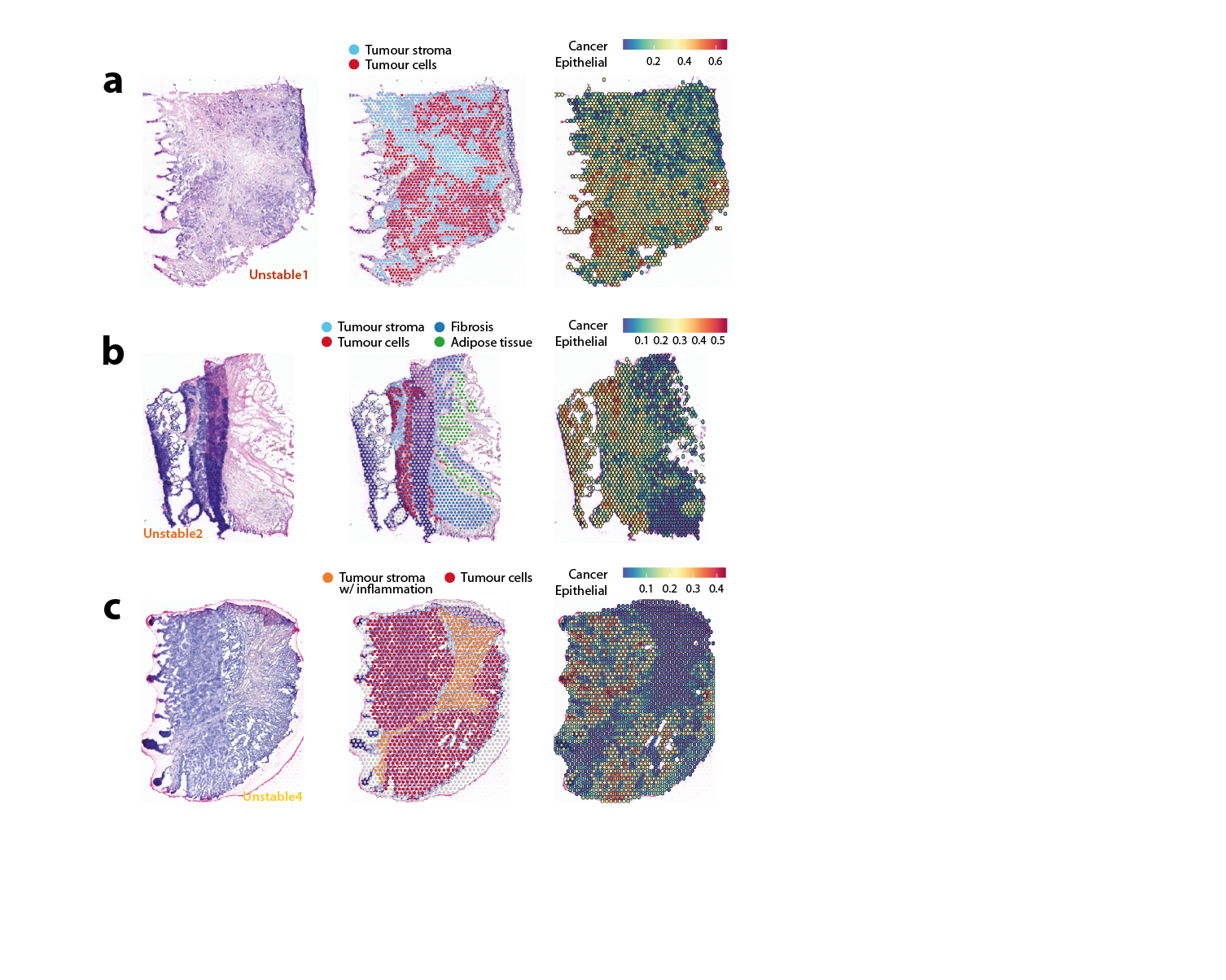

***Supplementary Fig. 7:*** *Hematoxylin & eosin staining (left), pathologist annotations (centre) and per-spot deconvolution-based cancer epithelial score (right) for Unstable samples Unstable1, Unstable2, and Unstable4.*

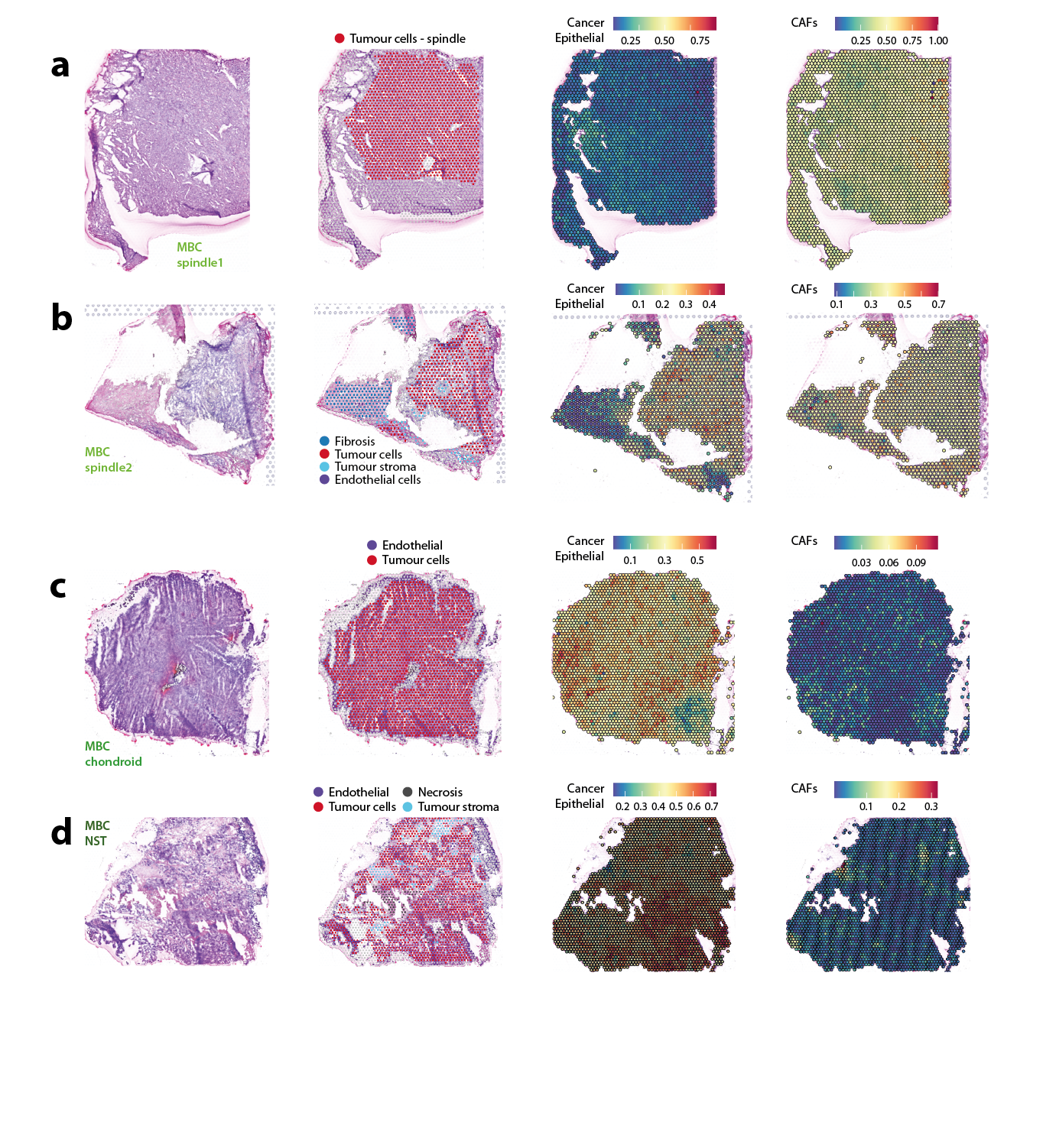

***Supplementary Fig. 8:*** *Hematoxylin & eosin staining (left), pathologist annotations (centre left), as well as per-spot deconvolution-based cancer epithelial score (centre right) and CAF score (right), for all MpBC samples.*

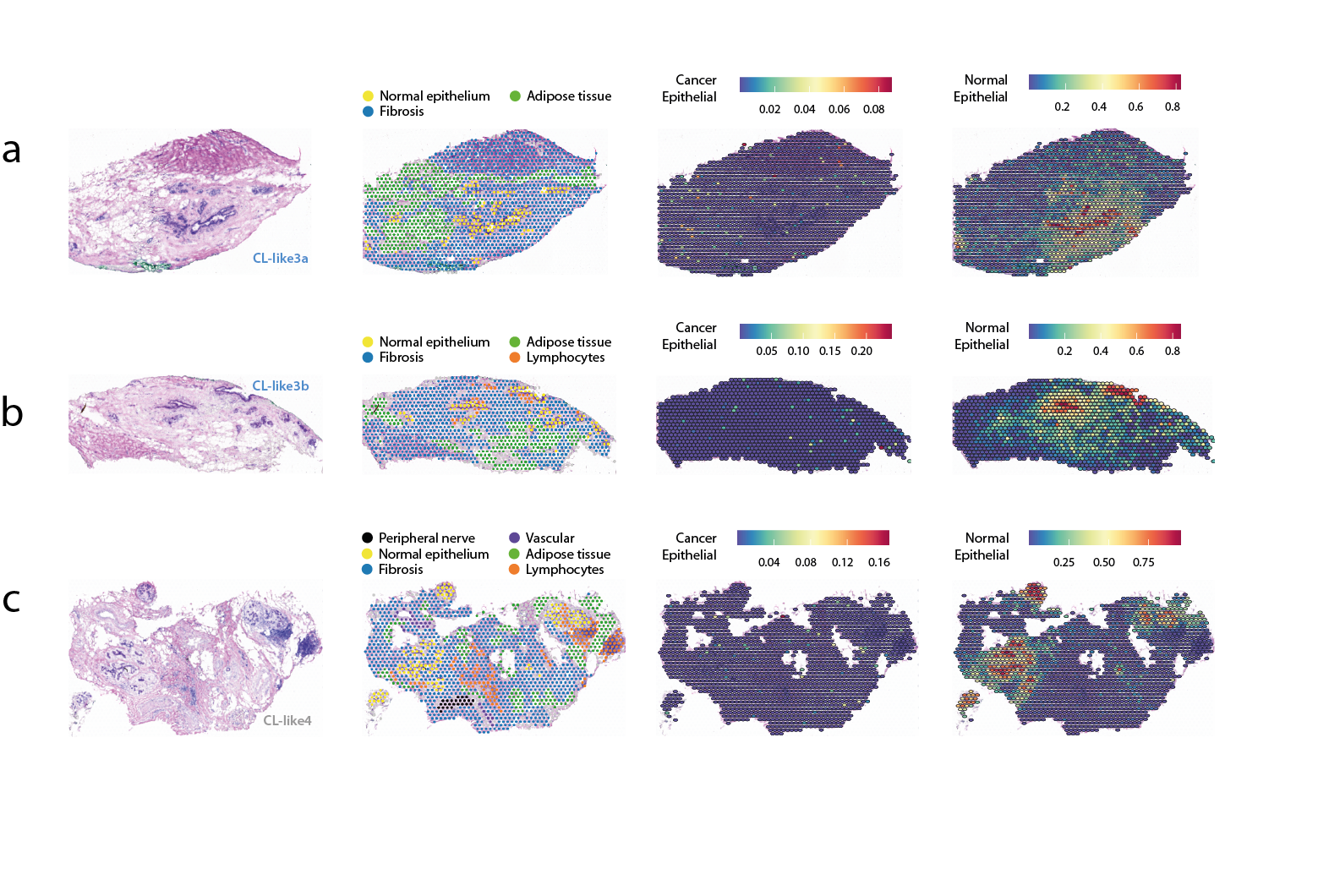

***Supplementary Fig. 9:*** *Hematoxylin & eosin staining (left), pathologist annotations (centre left), as well as per-spot deconvolution-based cancer epithelial score (centre right) and normal epithelial score (right), for all tumour-free samples used as normal references. Samples CL-like3a and CL-like3b are different tumour-free slides cut from the same tumour piece.*

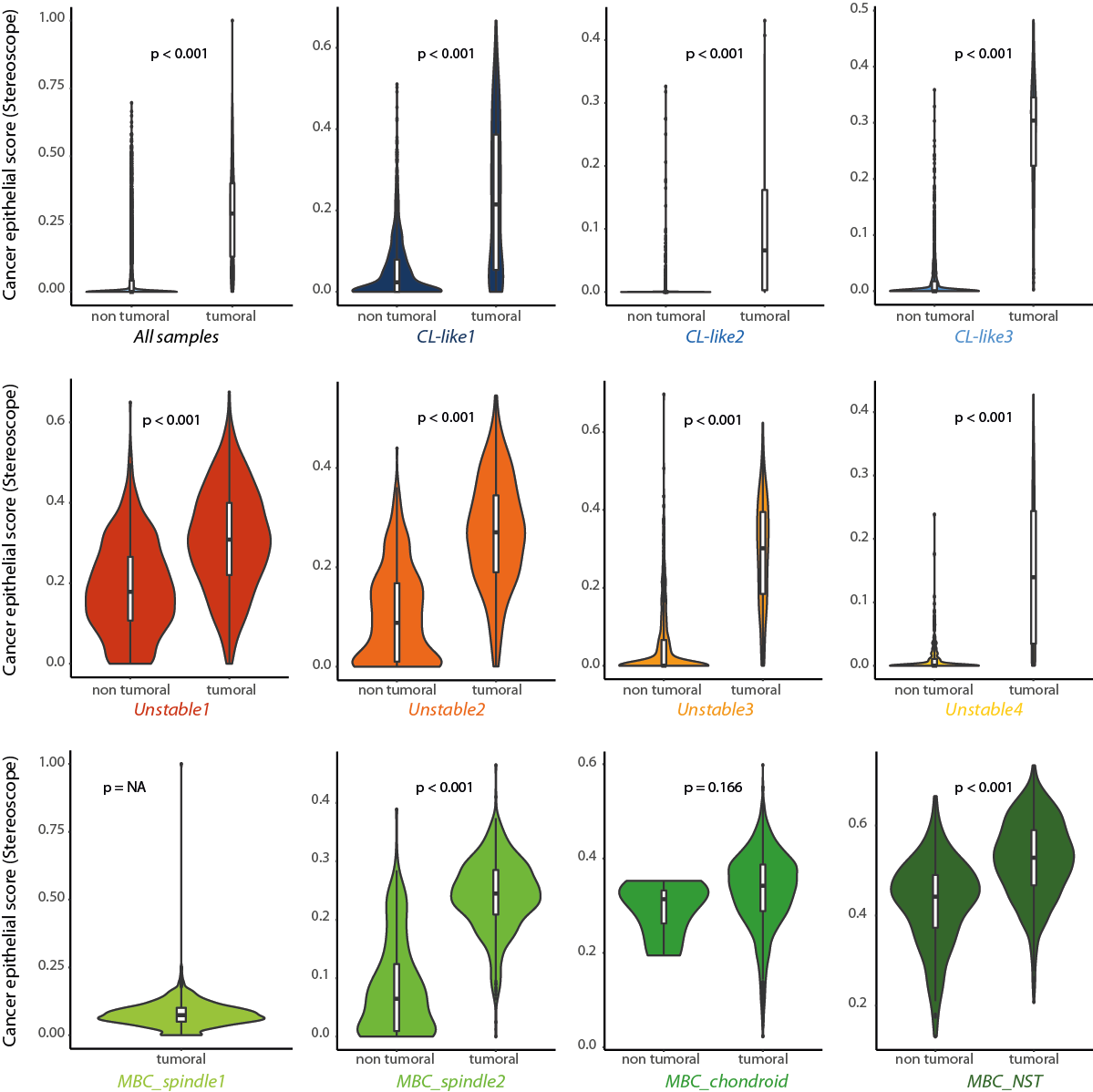

***Supplementary Fig. 10:*** *Per-spot deconvolution-based cancer epithelial scores for SpaT spots annotated as tumour and non-tumour, in all tumour slides pooled (top left) and individual slides.*

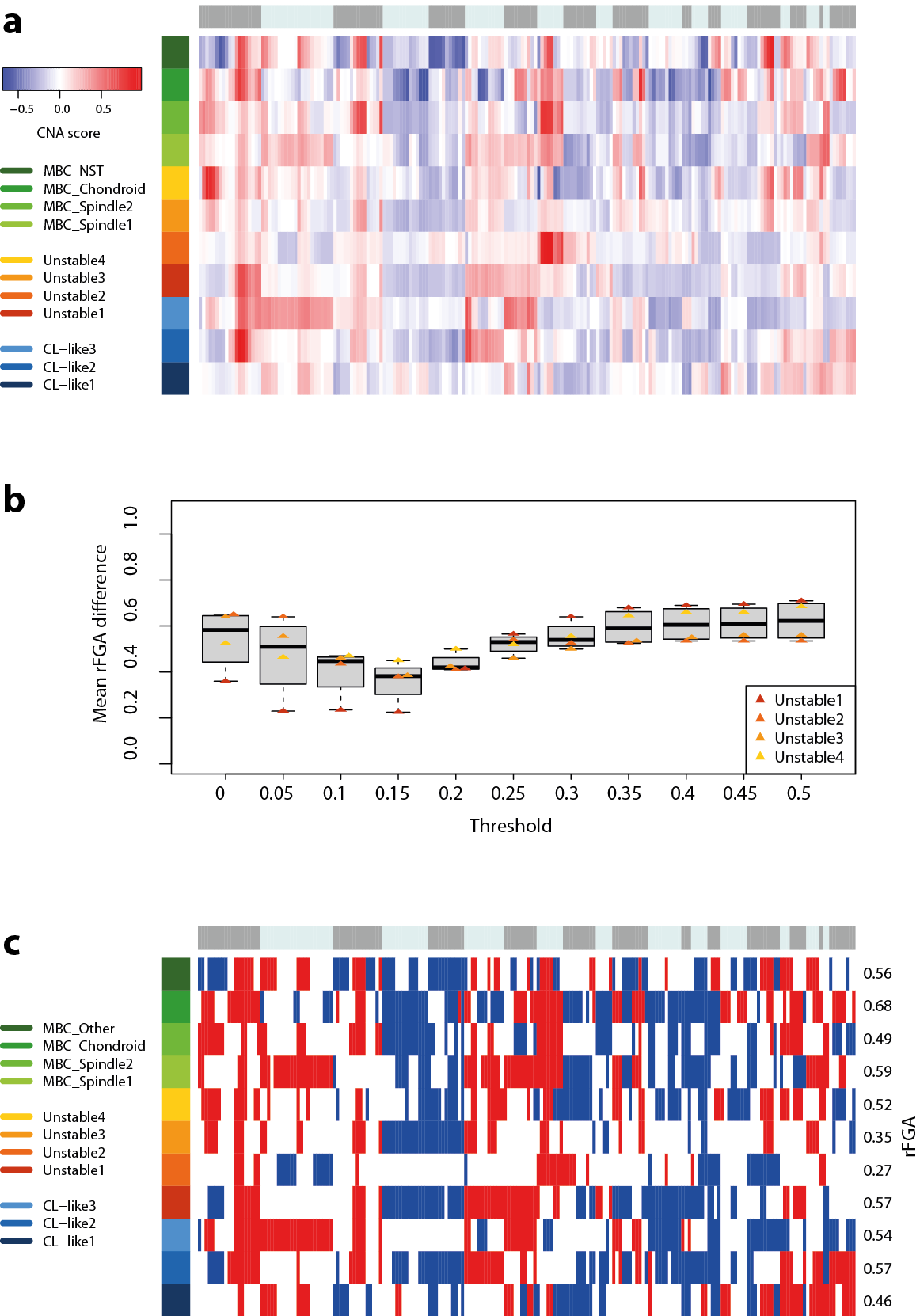

***Supplementary Fig. 11:*** *a) SpaT-derived CNA scores averaged per cytoband across all tumour slides. Cytobands as columns, samples as rows. Chromosome locations are indicated by alternating grey and light blue bars on top of the heatmap. b) difference in relative FGA (rFGA, based on the deviation from a sample’s average ploidy rather than from “normal” diploidy) between bulk and SpaT data for Unstable samples, according to the threshold used to define CNAs in CNA scores from SpaT data. 0.15 was selected as the most adequate threshold value. c) Gain/loss profiles defined using a 0.15 threshold on SpaT-derived CNA scores. Blue: loss (CNA score < -0.15); red: gain (CAN score > 0.15).*

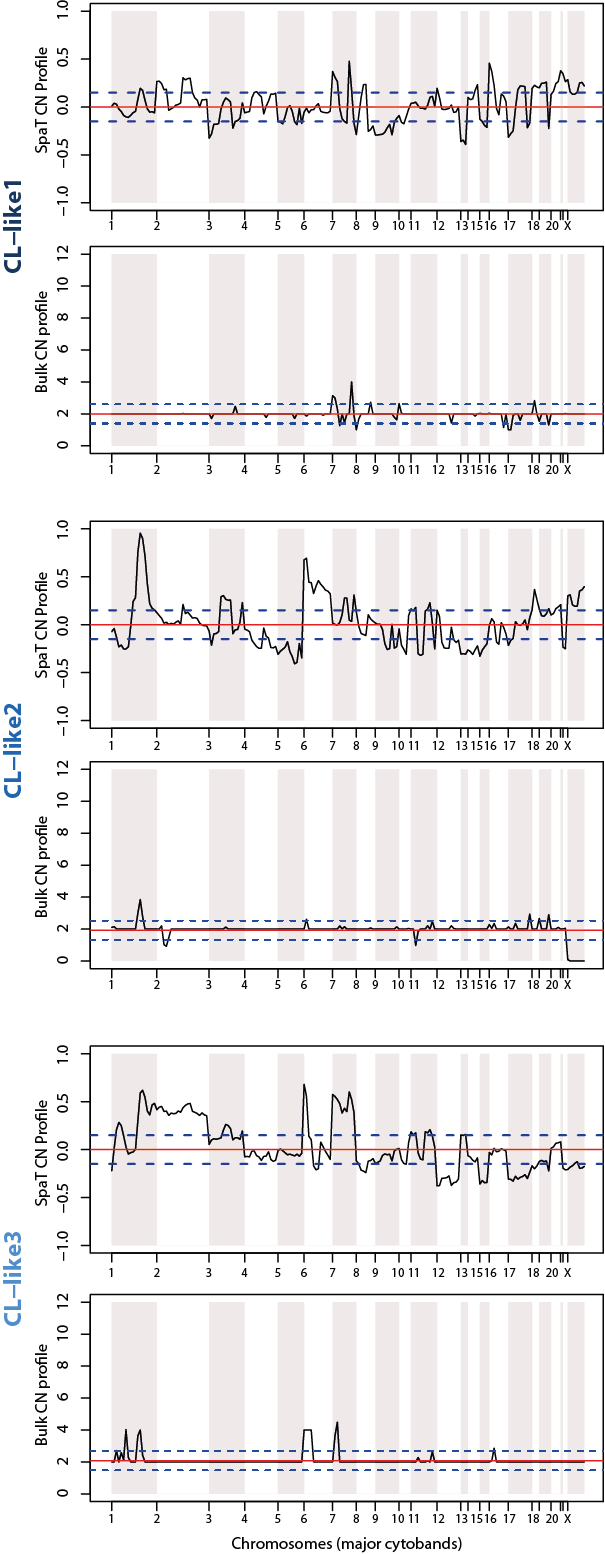
***Supplementary Fig. 12:*** *Overlay of SpaT-derived CNA scores averaged per cytoband (top) and bulk-derived total copy number profiles (bottom), for CL-like samples. The horizontal red line indicates the baseline for cytobands without copy number changes. The dashed horizontal blue lines indicate the thresholds used to consider whether a cytoband is gained (above the top blue line) or lost (below the bottom blue line). Chromosome locations are indicated by alternating white and grey backgrounds and specified on the x axis.*

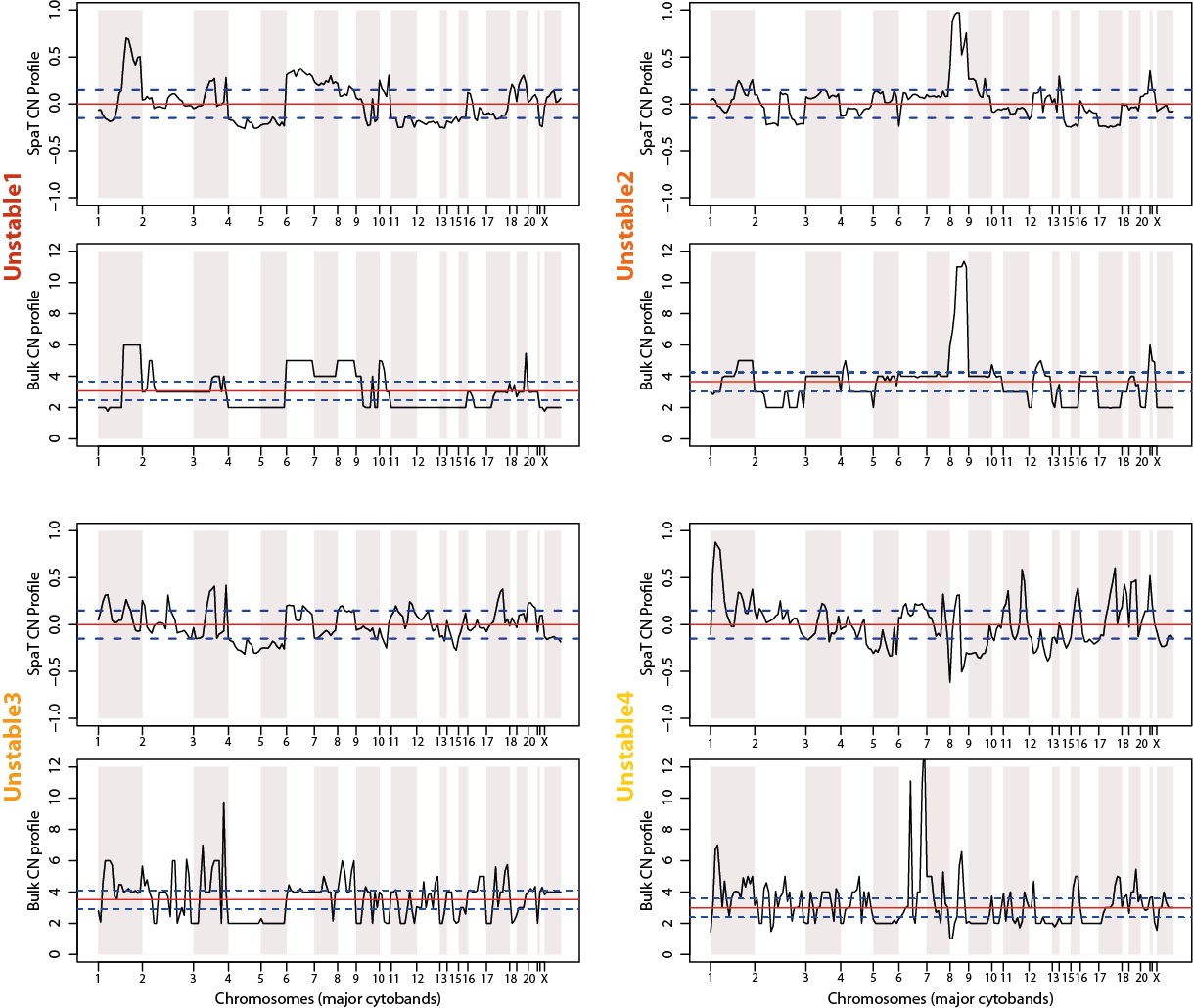

***Supplementary Fig. 13:*** *Overlay of SpaT-derived CNA scores averaged per cytoband (top) and bulk-derived total copy number profiles (bottom), for Unstable samples. The horizontal red line indicates the baseline for cytobands without copy number changes. The dashed horizontal blue lines indicate the thresholds used to consider whether a cytoband is gained (above the top blue line) or lost (below the bottom blue line). Chromosome locations are indicated by alternating white and grey backgrounds and specified on the x axis.*

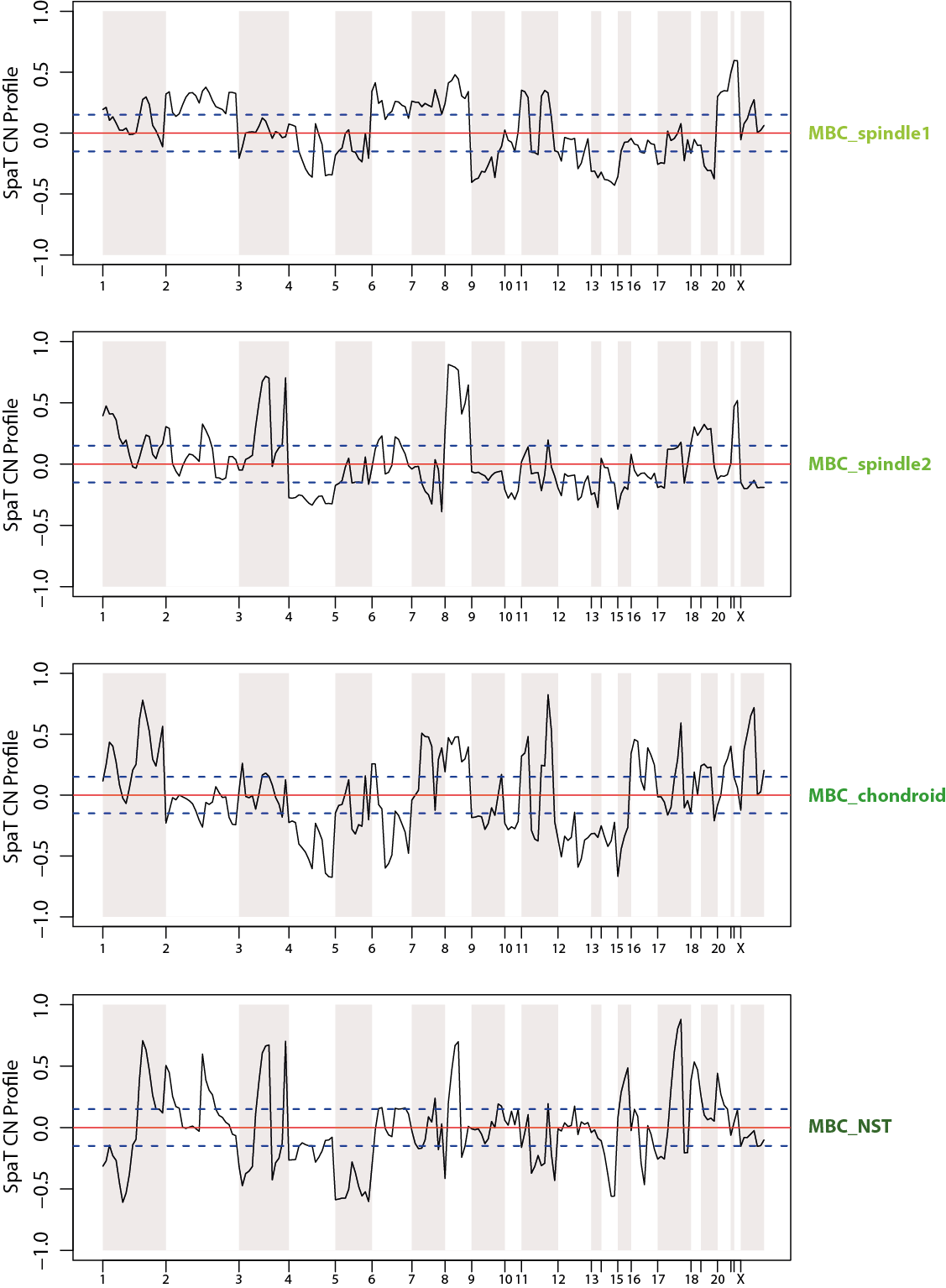

***Supplementary Fig. 14:*** *SpaT-derived CNA scores averaged per cytoband for MpBC samples. The horizontal red line indicates the baseline for cytobands without copy number changes. The dashed horizontal blue lines indicate the thresholds used to consider whether a cytoband is gained (above the top blue line) or lost (below the bottom blue line). Chromosome locations are indicated by alternating white and grey backgrounds and specified on the x axis.*

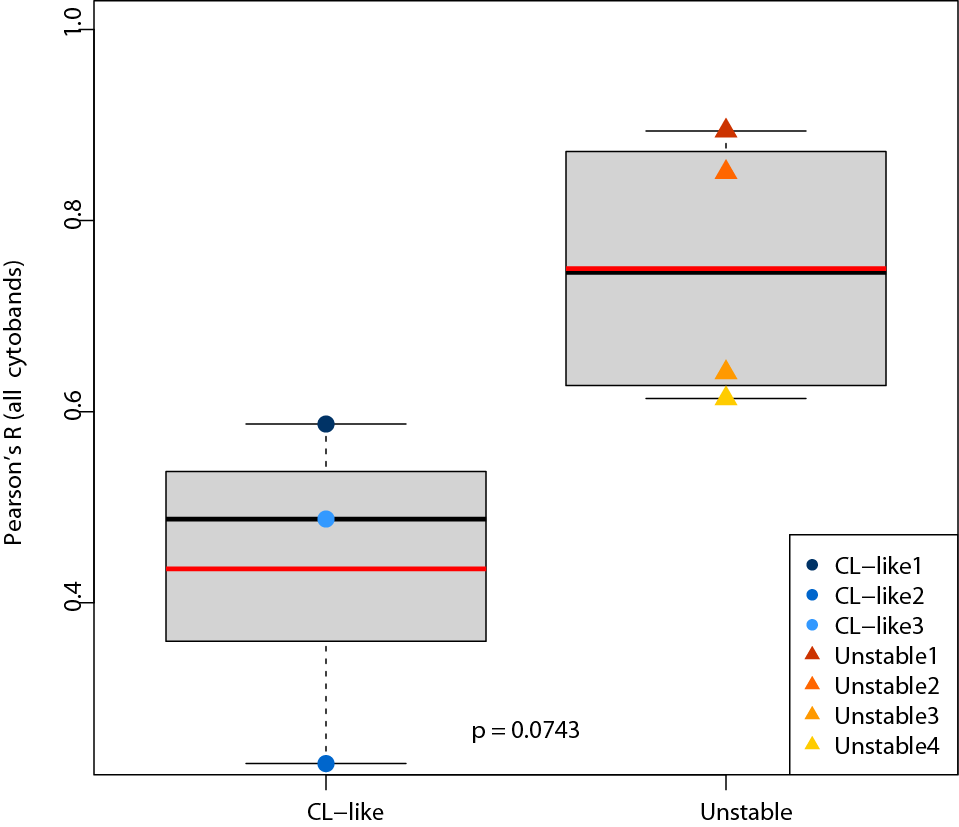

***Supplementary Fig. 15:*** *Pearson correlation between bulk logR and SpaT-derived average CNA scores, average per major cytoband, in CL-like and Unstable samples. All cytobands were included in the correlation calculation.*

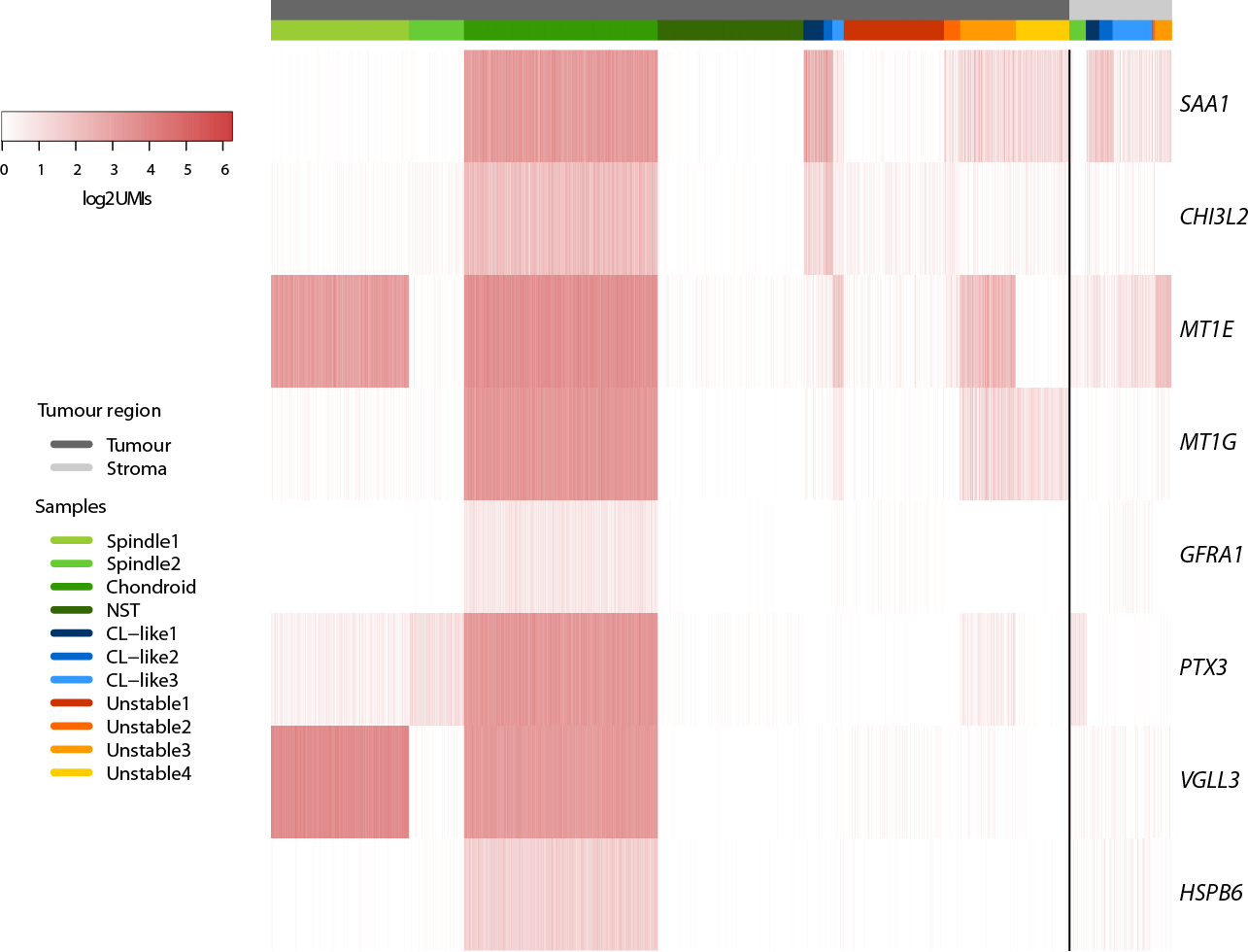

***Supplementary Fig. 16:*** *Expression heatmap for the newly identified chondroid MpBC markers. Expression is reported in log2 number of UMIs per spot, ranging from white (no expression) to red (high expression), and was analysed in both tumour spots (left, black) and stromal spots (right, grey).*

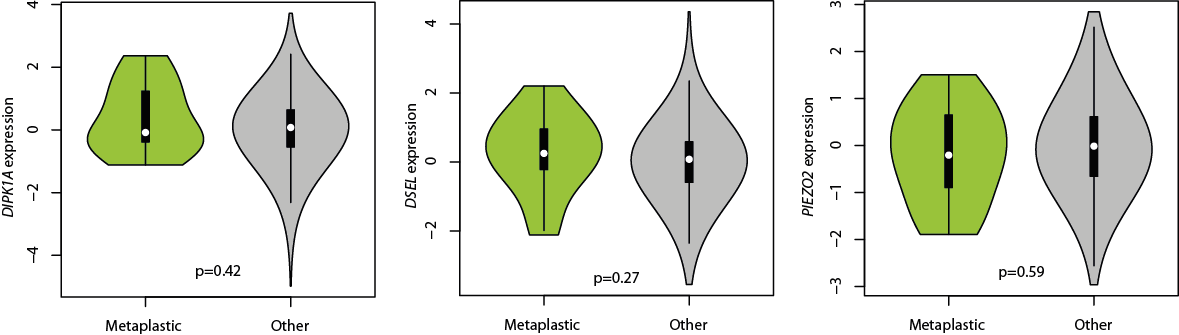

***Supplementary Fig. 17:*** *Expression of spindle cell MpBC candidate genes DIPK1A, DSEL, and PIEZO2, which could not be validated in an external cohort of 1100 samples analysed per bulk RNA-seq, stratified by tumour metaplastic status (14 metaplastic tumours in total). All p-values given by Wilcoxon rank-sum tests.*

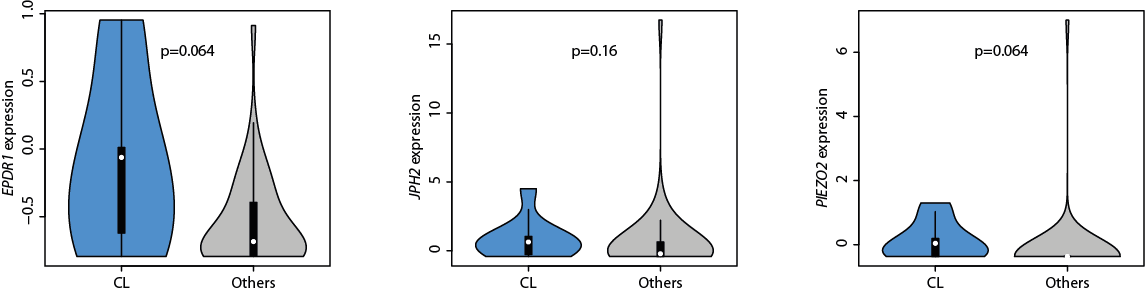

***Supplementary Fig. 18:*** *Expression of spindle cell MpBC candidate genes EPDR1, JPH2, and PIEZO2, which could not be validated in an external cohort of 51 samples analysed per bulk RNA-seq, stratified by tumour Claudin-low status (9 CL tumours in total). All p-values given by Wilcoxon rank-sum tests.*

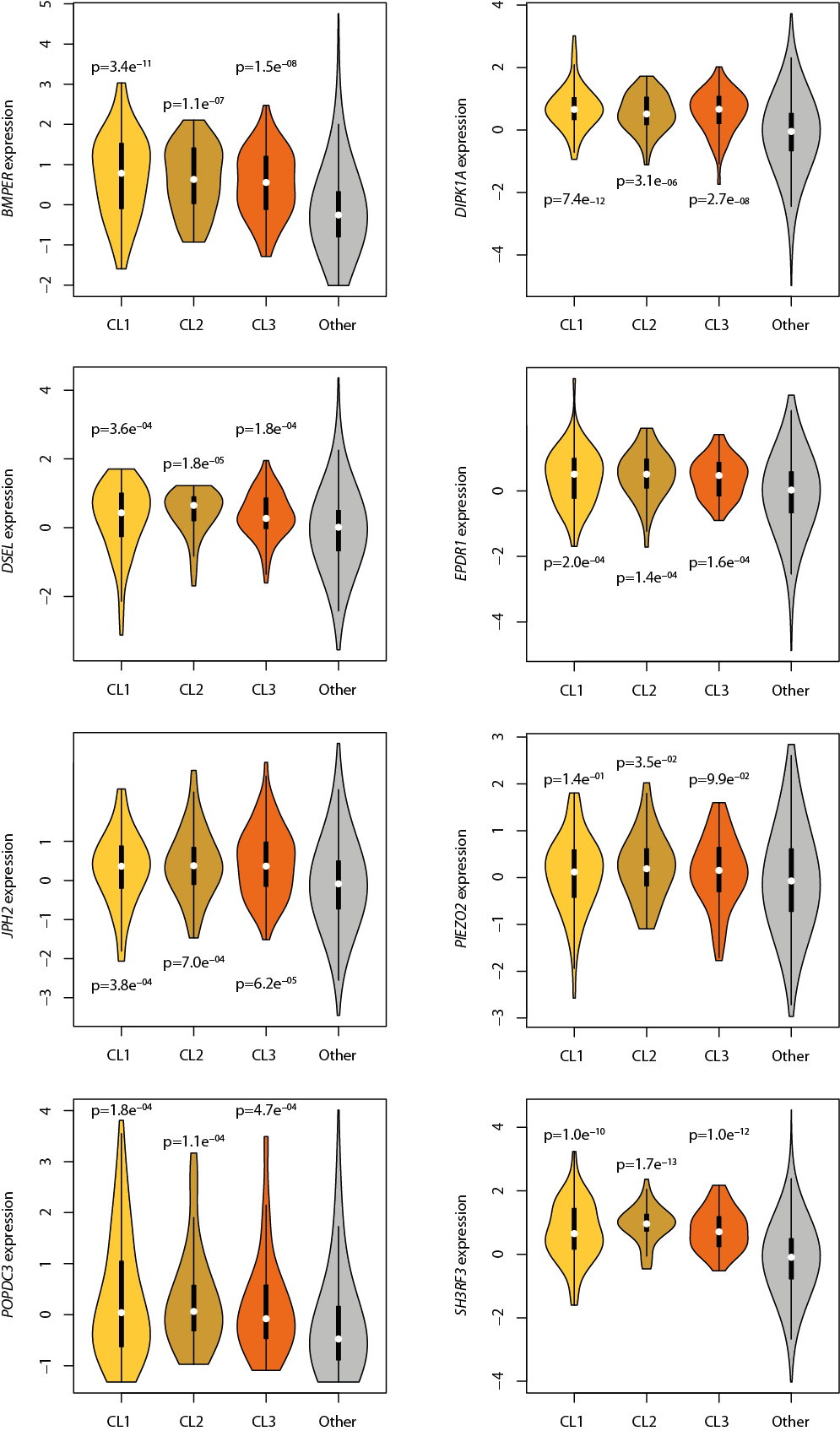

***Supplementary Fig. 19:*** *Expression of spindle cell MpBC marker candidate genes BMPER, DIPK1A, DSEL, EPDR1, JPH2, PIEZO2, POPDC3 and SH3RF3 in an external cohort of 1100 samples analysed per bulk RNA-seq, stratified by Claudin-low status (69 CL1, 42 CL2, 57 CL3 and 940 Other). All p-values given by Wilcoxon rank-sum tests.*

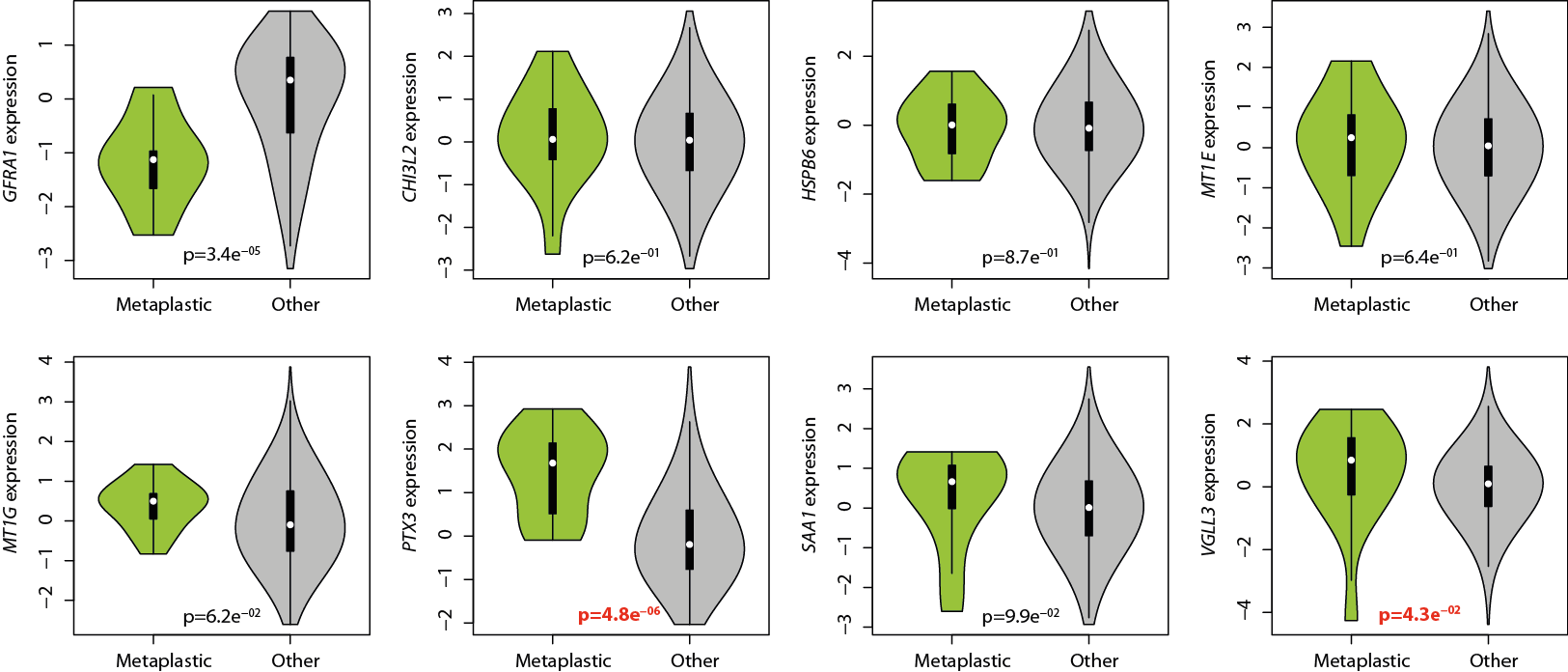

***Supplementary Fig. 20:*** *Expression of chondroid marker candidate genes GFRA1, CHI3L2, HSPB6, MT1E, MT1G, PTX3, SAA1 and VGLL3 in an external cohort of 1100 samples analysed per bulk RNA-seq, stratified by tumour metaplastic status (14 metaplastic tumours in total). All p-values given by Wilcoxon rank-sum tests; validated genes with overexpression and p-values < 0.05 are highlighted in bold red.*

| **Capture area ID** | **Visium Slide ID** | **Sequencing Batch** | **Sample ID** | **MyPROBE Tumour ID** | **Bulk FGA (%)** | **Tumour Cells** | **Spots Under Tissue** | **Fraction Reads Under Tissue** | **Mean Reads Per Spot** | **Median Genes Per Spot** | **Median UMIs Per Spot** | **Sequencing Saturation** |
| --- | --- | --- | --- | --- | --- | --- | --- | --- | --- | --- | --- | --- |
| M1 | 1 | 1 | CL-like3 | CLB-52 | 4.19 | NO | 1,689 | 89.4 | 213,729 | 909 | 1,424 | 98.0 |
| M2 | 1 | 1 | CL-like1 | CLB-17 | 9.12 | YES | 1,848 | 87.0 | 171,777 | 2,398 | 5,083 | 91.7 |
| M3 | 1 | 1 | Unstable1 | CLB-23 | 78.72 | YES | 2,417 | 83.8 | 92,937 | 2,543 | 5,634 | 84.6 |
| M4 | 1 | 1 | Unstable2 | CLB-37 | 76.28 | YES | 1,564 | 65.4 | 144,123 | 2,266 | 5,548 | 85.1 |
| M5 | 2 | 1 | Unstable3 | CLB-51 | 80.93 | YES | 2,104 | 98.0 | 68,204 | 2,078 | 3,972 | 84.8 |
| M6 | 2 | 1 | Unstable4 | CLB-14 | 80.62 | YES | 1,731 | 90.7 | 121,664 | 4,595 | 21,851 | 76.8 |
| M7 | 2 | 1 | CL-like4 | CLB-11 | 4.72 | NO | 1,596 | 91.8 | 129,152 | 1,254 | 2,369 | 94.4 |
| M8 | 2 | 1 | CL-like2 | CLB-74 | 9.96 | YES | 1,055 | 87.2 | 230,372 | 2,226 | 4,950 | 93.1 |
| M9 | 3 | 1 | CL-like3 | CLB-52 | 4.19 | NO | 1,258 | 85.2 | 133,505 | 745 | 1,205 | 95.1 |
| M10 | 3 | 1 | CL-like4 | CLB-11 | 4.72 | NO | 370 | 76.0 | 181,917 | 1,383 | 2,616 | 94.7 |
| M11 | 3 | 1 | CL-like3 | CLB-52 | 4.19 | YES | 1,405 | 90.6 | 164,993 | 2,799 | 7,446 | 87.7 |
| M13 | 4 | 2 | MBC_spindle1 | NA | NA | YES | 2,657 | 82.1 | 153,132 | 4,716 | 15,124 | 75.7 |
| M14 | 4 | 2 | MBC_spindle2 | NA | NA | YES | 1,295 | 73.9 | 144,904 | 4,146 | 11,126 | 79.6 |
| M15 | 4 | 2 | MBC_chondroid | NA | NA | YES | 3,037 | 78.8 | 62,673 | 5,066 | 19,634 | 50.1 |
| M16 | 4 | 2 | MBC_NST | NA | NA | YES | 2,700 | 80.8 | 79,589 | 4,266 | 2,334 | 66.9 |

***Supplementary Table 1****: Spatial transcriptomics sample information.*
